# Use of the *p-value* as a size-dependent function to address practical differences when analyzing large datasets

**DOI:** 10.1101/2019.12.17.878405

**Authors:** Estibaliz Gómez-de-Mariscal, Vanesa Guerrero, Alexandra Sneider, Hasini Jayatilaka, Jude M. Phillip, Denis Wirtz, Arrate Muñoz-Barrutia

## Abstract

Biomedical research has come to rely on *p-values* as a deterministic measure for data-driven decision making. In the largely extended null-hypothesis significance testing (NHST) for identifying statistically significant differences among groups of observations, a single *p-value* computed from sample data is routinely compared with a threshold, commonly set to 0.05, to assess the evidence against the hypothesis of having non-significant differences among groups, or the null hypothesis. Because the estimated *p-value* tends to decrease when the sample size is increased, applying this methodology to large datasets results in the rejection of the null hypothesis, making it not directly applicable in this specific situation. Herein, we propose a systematic and easy-to-follow method to detect differences based on the dependence of the *p-value* on the sample size. The proposed method introduces new descriptive parameters that overcome the effect of the size in the *p-value* interpretation in the framework of large datasets, reducing the uncertainty in the decision about the existence of biological/clinical differences between the compared experiments. This methodology enables both the graphical and quantitative characterization of the differences between the compared experiments guiding the researchers in the decision process. An in-depth study of the proposed methodology is carried out using both simulated and experimentally obtained data. Simulations show that under controlled data, our assumptions on the *p-value* dependence on the sample size holds. The results of our analysis in the experimental datasets reflect the large scope of this approach and its interpretability in terms of common decision-making and data characterization tasks. For both simulated and real data, the obtained results are robust to sampling variations within the dataset.

## INTRODUCTION

The ability to acquire, store and disseminate large amounts of data is constantly improving in life-science laboratories. Having such big datasets available for multiple kinds of analysis supports the proliferation of many different analytic applications. Nonetheless, it has also emphasized the challenges that classical statistical techniques need to face when analyzing such types of datasets, often called Big Data Analysis. An extended practice in experimental life-science is the analysis of differences among experimental settings, which is usually determined through the mean values of the studied statistical variable in distinct groups. In order to decide whether statistically significant differences exist, null hypothesis significance testing (NHST) is usually performed. Namely, a formal hypothesis test is stated in which the no effect hypothesis, that is the equality of the mean values yielded by experimental datasets, is assessed thanks to the computation of a *p-value* on sample data. This value is then compared with the threshold 0.05 to decide whether or not the null hypothesis is rejected. When working with large datasets, the accuracy of the estimators of those mean values (or other parameters) improves, Fig 1 and 2 in the Materials and Methods. While NHST nearly always finds statistical differences among the group means in large datasets, the researchers usually aim to find out whether those differences are *interesting*, e.g. biologically or clinically relevant. Technically, the *p-value* depends on the size of the data being tested: the larger the sample size, the smaller the *p-value*. An easy understanding of the latter relies on the evidence in the data against the null hypothesis instead of the existence of interesting differences among groups (Greenland, 2019). The larger the data size, the larger the accuracy of the statistical test, and therefore, the larger the evidence against the null hypothesis. The latter is in high contrast with the recurrent misleading interpretation of the *p-value* as a “gold standard” for the identification of biologically or clinically relevant differences among experiments (Altman and Krzywinski, 2017; Amrhein et al., 2019; Greenland, 2019; Halsey et al., 2015; Nuzzo, 2014). In particular, when large sample are available, life-scientists could detect statistically significant evidence against the null hypothesis through a small enough *p-value*, even though there are no interesting differences from the practical point of view. Even more, the *p-value* is itself a random variable that depends on the sample data used to estimate it; and, therefore, has a sampling distribution that is intrinsically determined by the noise in the data. A straightforward example is as follows: the *p-value* has a uniform distribution 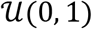 under the null hypothesis, which is rejected 5% of the times when a significance threshold of 5% is being used. Hence in 5% of the cases, a single computation of the *p-value* could lead to the contradictory conclusion that there exist statistically significant differences among two groups identically distributed (Bruns and Ioannidis, 2016). Similar to the examples in (Altman and Krzywinski, 2017; Halsey et al., 2015), Fig. 5 in the Materials and Methods further illustrates the described behavior.

**Fig. 1.**
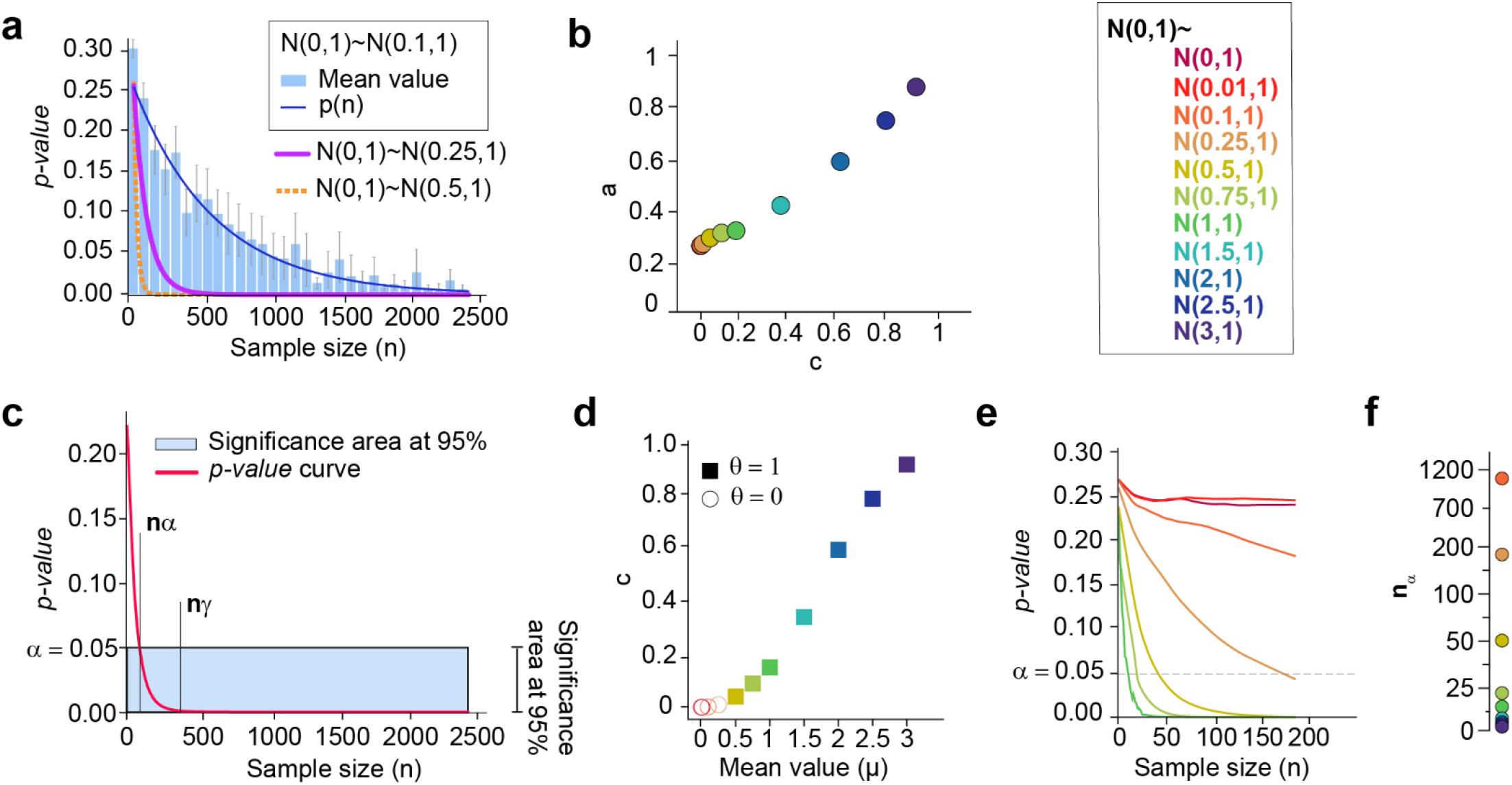
Estimation of the *p-value* as a function of the size (*p*(*n*)) enables the correct discrimination between conditions. **a)** The *p-value* is a variable that depends on the sample size and can be modelled as an exponential function (*p*(*n*) = *ae^−cn^*, Eq. 1). For each pair of normal distributions being compared, two subsets of size *n* are obtained by sampling from the corresponding normal distribution. Then, these datasets are compared using the Mann-Whitney statistical test and the *p-value* obtained is stored. The procedure is repeated many times for each size *n*. The blue bars with the standard error of the mean (SEM), show the distribution of all the *p-values* obtained at each size *n* when two normal distributions of mean 0 and 0.1, and standard deviation 1 are compared. The blue curve shows the corresponding exponential fit. The magenta and yellow curves represent the resulting *p*(*n*) function when a normal distribution of mean 0 and standard deviation 1 is compared with a normal distribution of the same standard deviation and mean 0.25 and 0.5, respectively; **b)** The decay of *p*(*n*) (parameters *a* and *c* of the exponential fit) increases with the mean value of the normal distribution being compared with *N*(0, 1). The larger the distances between the means of the distributions, the higher the decay of the exponential function (Table 1 in Materials and Methods). **c)** Comparison of *p*(*n*) (red curve) and significance area at 95% (blue area). If the area under the red curve is smaller than the blue area, then there is a strong statistical significance. The parameter *n_α_* measures the minimum data size needed to find statistical significance. The parameter *n_γ_* measures the convergence of *p*(*n*): *p*(*n* = *n_γ_*) ≈ 0. The binary decision index *θ* indicates whether the area under *p*(*n*) from 0 to *n_γ_* is larger than the area under the *α*-level (blue box) in the same range; **d)** The faster the decay of *p*(*n*), the stronger the statistical significance of the tested null hypothesis. For *γ* = 5*e*^−06^, *θ_α,γ_* = 1 whenever the mean value of the normal distribution compared with *N*(0, 1) is larger than 0.5 (Table 1 in Materials and Methods). **e)** The empirical estimation of *p*(*n*) with small datasets enables the detection of the most extreme cases: those in which the null hypothesis can be accepted, and those in which it clearly cannot; **f)** The minimum data size needed to obtain statistical significance (*n_α_*) is inverse to the mean value of the normal distributions being compared.

**Figure 1.**
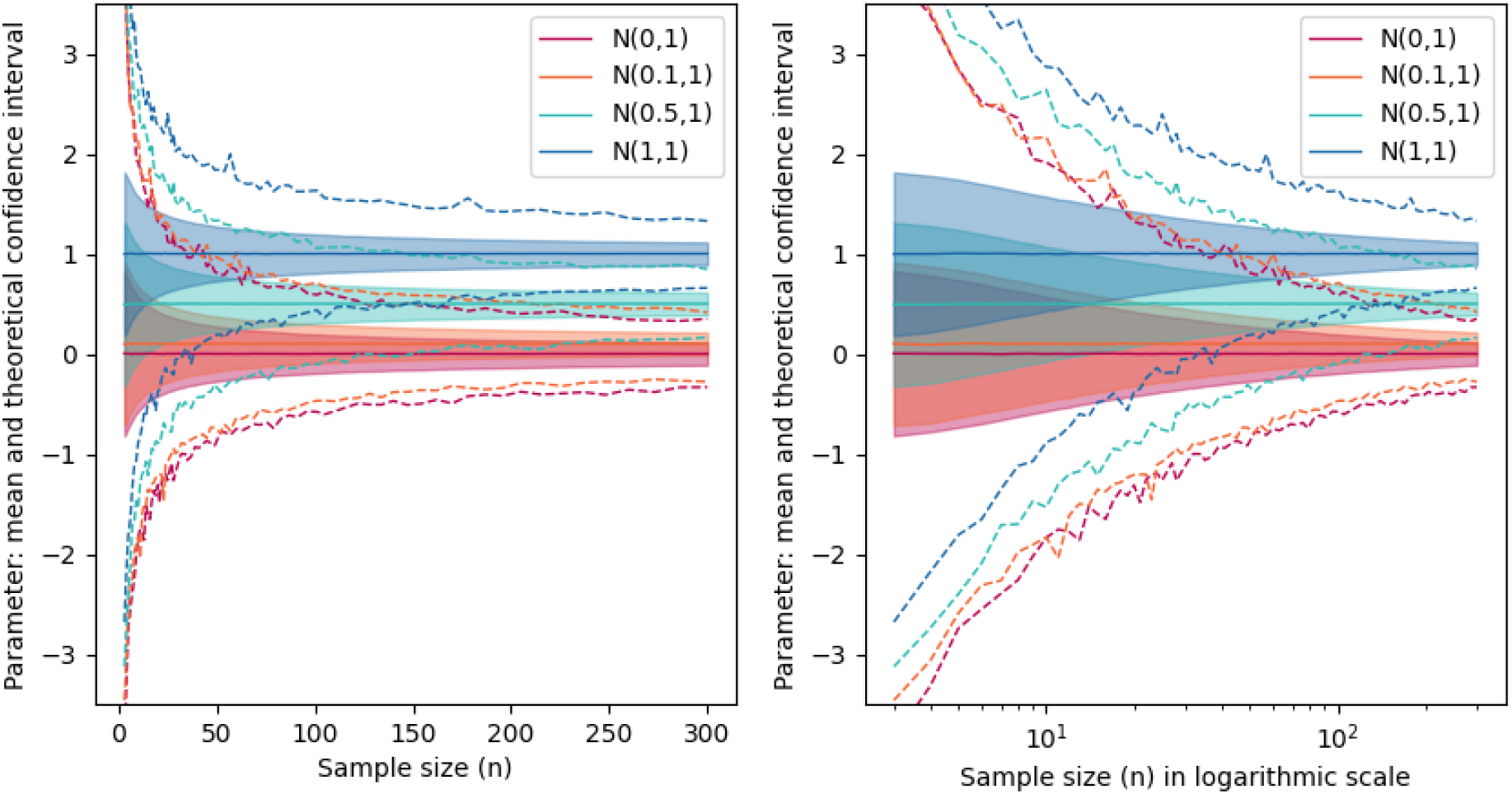
Estimated two-sided 95% confidence interval for the mean of different normal distributions with standard deviation of 1 and mean values of 0, 0.1, 0.5 or 1. For each fixed value of the sample size, 15000 confidence intervals for the mean of each normal distribution were calculated using the mathematical expression 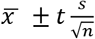, where 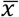 is the estimated mean value, *t* is the critical value of Student’s t-distribution for a 0.025 of significance, *s* is the estimated variance and *n* is the sample size (filled area). The dashed lines show the maximum and minimum values of the calculated confidence intervals obtained for each sample size. The information is shown both in linear and logarithmic scale.

**Fig. 2.**
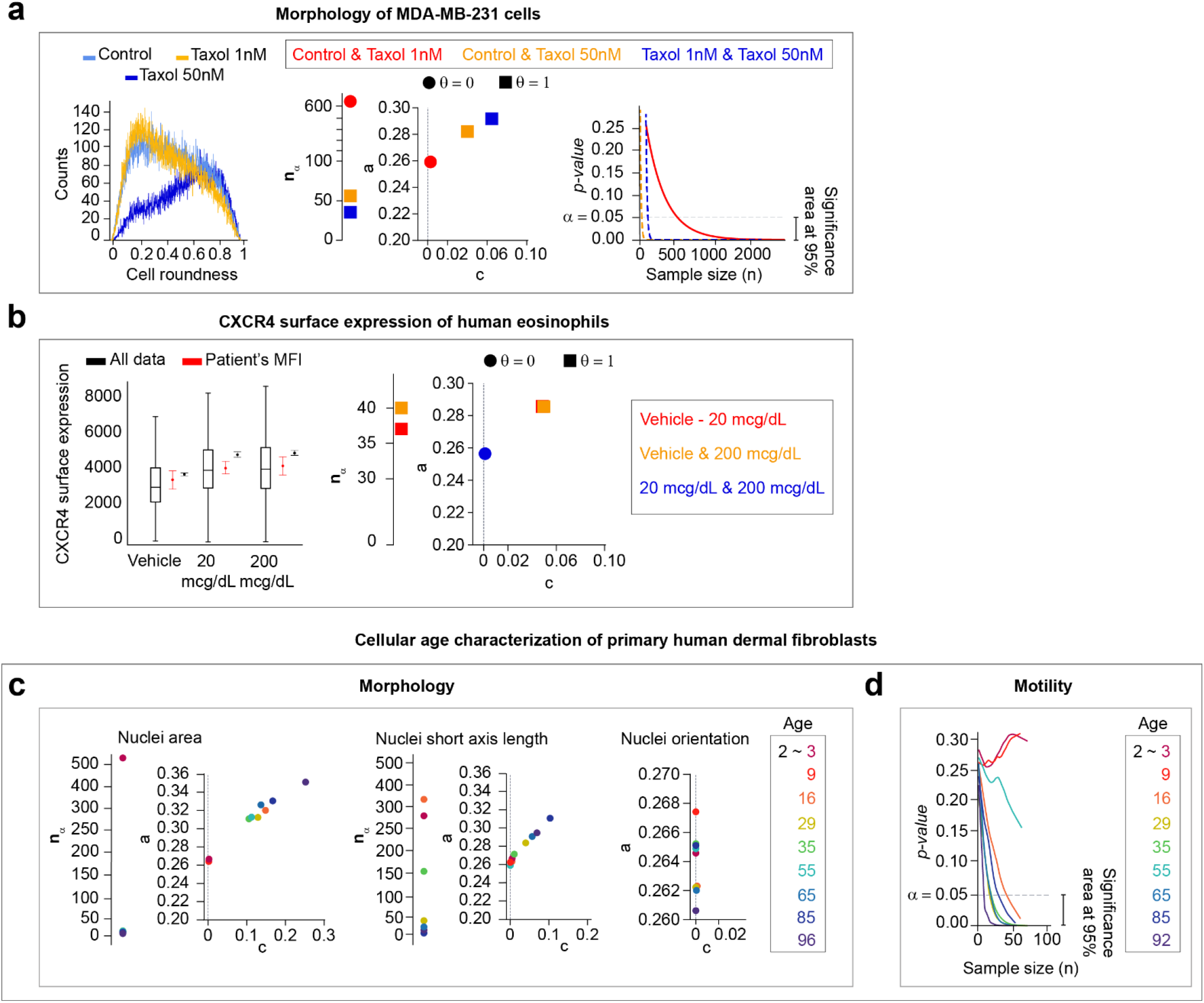
The function *p*(*n*) acts as a data descriptor and supports the experimental study of multiple conditions. **a)** Breast cancer cells (MDA-MB-231) were cultured in collagen and imaged under a microscope to determine if cells change shape when a chemotherapy drug (Taxol) is administered. Three different groups were compared: control (non-treated) cells, cells at 1 nM and at 50 nM Taxol. (Leftmost) The cell roundness distribution of control cells and cells treated at 1 nM Taxol have lower values than that of cells treated at 50 nM. (Right) The three groups were compared, the *p-values* were estimated and *p*(*n*) was fitted for each pair of compared groups. When Taxol at 50 nM is evaluated (blue and yellow dashed curves), *n_α_* is lower and the decay of *p*(*n*) is higher (*a* and *c* parameters in Eq. 1), i.e. it decreases much faster than the one corresponding comparison of control and Taxol at 1 nM (orange curve). **b)** Flow cytometry data was recorded to determine the transcriptional changes induced by the in vivo exposure of human eosinophils to glucocorticoids. (Left) The entire dataset has a wider range of values (black box-plots) and a smaller 95% confidence interval around the mean (black error-plots) than the distribution obtained when the median fluorescence intensity (MFI) is calculated by each of the 6 subjects (red error-plots). (Right) There is an increase of the surface expression of CXCR4 when human eosinophils are exposed to 20 or 200 mcg/dL of Methylprednisolone. Namely, the minimum size *n_α_* is low and the decision index *θ* = 1 when any of those conditions are compared with the vehicle condition. Note that the decay parameters *a* and *c* are almost the same in those two cases, so the markers co-localize (Supplementary Material). The minimum size *n_α_* when eosinophils are treated (blue circle) is not shown as it has infinite value. **c)** The morphology of 2 year-old human cells is compared with the morphology of 3, 9, 16, 29, 35, 45, 55, 65, 85 and 96 year-old human cells. For both, nuclei area and nuclei short axis measures, the minimum size *n_α_* and the decay *α* change proportionally with the age of the donor. The nuclei orientation does not characterize the age of the human donors for all the comparisons; the parameter *c* is null, and therefore, *p*(*n*) is constant. **d)** The analysis of a small dataset is enough to determine that the total diffusivity can characterize the cellular aging in humans. The total diffusivity of 2, 3 and 9 year-old human cells are equivalent, while it differs when compared to cells from older human donors.

**Figure 2.**
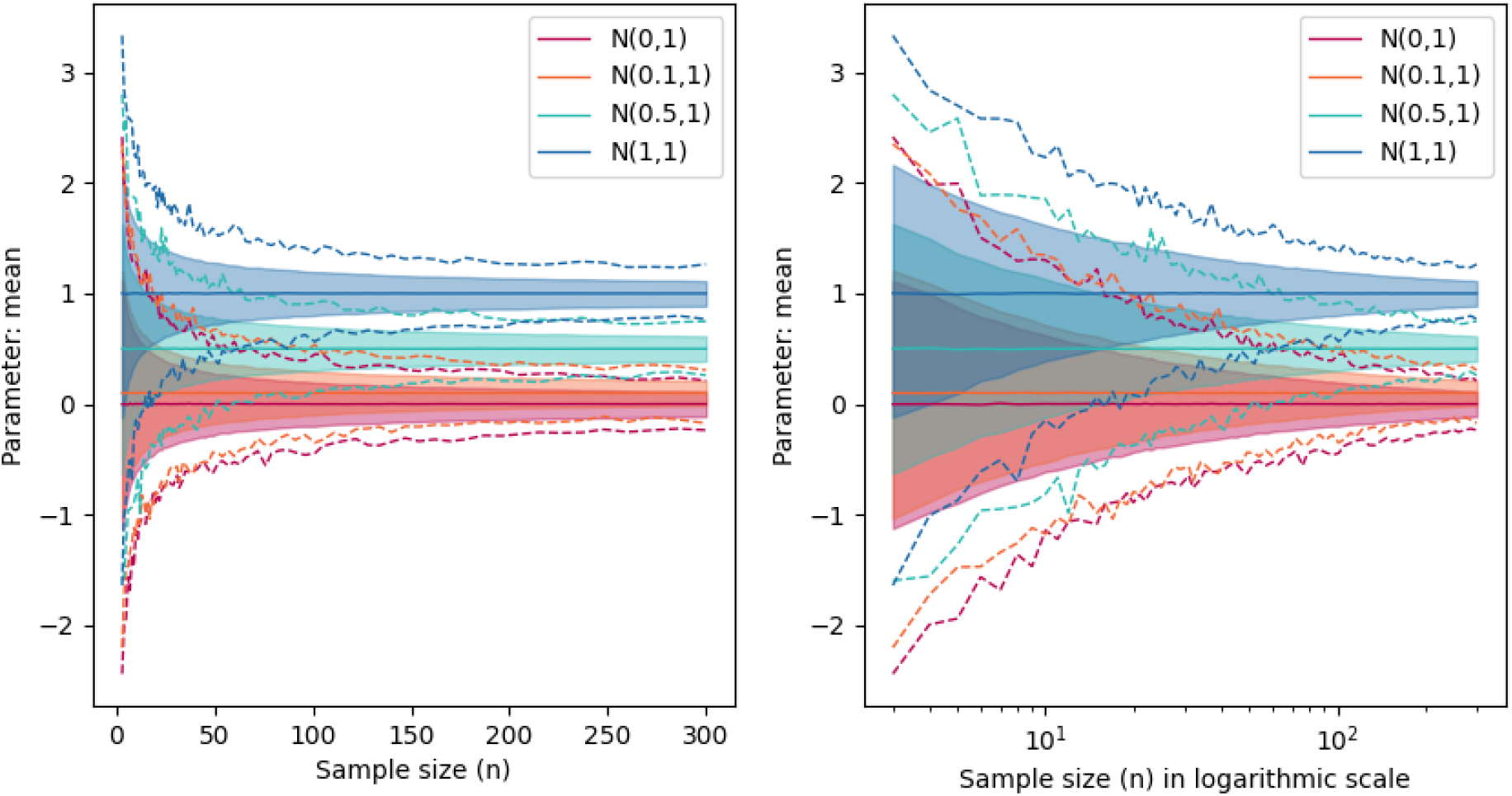
Bootstrapping estimation of two-sided 95% confidence interval for the mean of different normal distributions with standard deviation of 1 and mean values of 0, 0.1, 0.5 or 1. For each value of the sample size, we compute the mean of a simulated normal distribution 15000 times. The final confidence interval is obtained by clipping 95% of the values among the 15000 (filled area). The dashed lines show the maximum and minimum values of the sample mean value obtained for each sample size. The information is shown both in linear and logarithmic scale.

Despite this finding, there remain many situations for which the ‘dichotomy’ associated with the *p-value* is necessary for data-driven decision-making (Leek et al., 2017). Here, we present different approaches described in the literature. An alternative solution to facilitate the interpretability of the *p-value* is the use of Shannon information or S-values (*s* = −log_2_ *p*, where *p* is the *p-value*) (Greenland, 2019). The S-values express the self-information given by the datasets with respect to the null-hypothesis rather than a probability or evidence against it. However, the S-value, same as the *p-value*, depends on the sample size and the datasets used to compute a single realization of them. Therefore, used as single numbers, they have a limited capacity to inform about practical relevance of the differences among the compared groups (Greenland, 2019). Other approaches analyze the distribution of empirically estimated *p-values*, also known as *p-curve*, (Simonsohn et al., 2014), which does not take into account the effect of the sample size. Computing the *p-curve* for large datasets will result in a high frequency of *p-values* around zero regardless the differences among the compared groups. There are also works that focus on the sample-size-dependence and the sensitivity of the *p-value* to this size, commonly denoted by *n* (Lin et al., 2013). The authors provide a detailed description of the drawbacks of NHST applied to large datasets and they suggest the use of confidence intervals and effect sizes as alternative measures. However, to the best of our knowledge, there are no methods that exploit the exponential behavior of the *p-value* using easily interpretable parameters to assess the existence of interesting differences from the biological or clinical perspective. In this work, an exponential model is fitted, which approximates the *p-value* for a continuum of sample sizes, in the context of finding statistically significant evidence against or in favor of differences among the mean values of an observed variable in two or more groups. Namely, we aim to answer the question of when we can solidly assert that bona fide differences exist from the practitioner’s perspective rather than just statistically significant. This paper presents an easily interpretable tool to support biomedical researchers in their statistical analyses.

Our method is based on assuming an exponential relationship between the sample size *n* and the *p-value*. We compute the Mann-Whitney U statistical test to find statistically significant differences between two or more groups. Then, for a given sample size *n* its *p-value*, *p*(*n*) is approximated using:

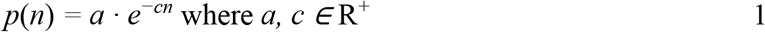

The values of *a* and *c* are found to minimize the squared differences between a set of observed *p-values* for each of the samples of size *n*, which are obtained using Monte Carlo cross-validation (MCCV) (Xu and Liang, 2001), and its estimation using the exponential approach. The parameters *a* and *c* are associated with the dependency of the *p-value* on the sample size and quantitatively measure the relationship between the two or more groups being compared.

In Fig. 1a, different randomly generated normal distributions are compared using the Mann-Whitney U statistical test (Mann and Whitney, 1947) to illustrate the decrease of the function *p*(*n*) with the sample size. The use of the Student’s *t*-test was avoided as it is well known that the *p-value* associated to the *t*-statistic has an exponential decay (Student, 1908). Technical details about the convergence of the function *p*(*n*) and evidence about Eq. 1 holding for any statistical test are given in the Materials and Methods.

Note that the *p-value* curve, the function *p*(*n*) in Eq. 1, is used to compare pairs of experimental conditions; therefore, *p*(*n*) is computed as the exponential fit of the *p-values* computed on multiple samples of different sizes extracted from the large dataset at hand. Hence, the parameters *a* and *c* in Eq. 1 correspond to those defining the exponential fit *p*(*n*). We use MCCV as the sampling strategy: two subsets of size *n* (one from each of the groups to be compared) are randomly sampled and compared with a statistical test for different values of the sample size *n*. The resulting *p-value* is stored and the procedure is repeated many times. At the end of the procedure, a large set of *n*-dependent *p-values* is obtained and the exponential function in Eq. 1 can be fit.

Similar to any exponential function, *p*(*n*) converges to zero. The faster the function converges, the stronger is the evidence against the null hypothesis. In other words, a fast decay implies finding statistically significant differences between the groups at small sample sizes, i.e. differences appear early. When normal distributions of standard deviation one and mean value in the range [0, 3] are compared, we see that the higher the difference among the mean values of each normal distribution, the faster the decay of the exponential function *p*(*n*), as expected (Fig. 1a). We observe that the parameters *a* and *c* (Eq. 1) increase proportionally with the mean value of the distribution compared with *N*(0, 1) (Fig. 1b). Thus, and enable the spatial representation of each normal distribution with respect to *N*(0, 1). These parameters can simplify the identification of the existence of interesting biological differences. Indeed, they can measure how far from each other the distributions of the groups being compared are (Fig. 1b).

With this new idea in mind, a robust decision index, *θ_α,γ_*, can be mathematically defined (Eq. 10 in Materials and Methods) which depends on the significance level *α* and a regularization parameter *γ* related to the convergence of the exponential fitted function. Note that subscripts *α* (statistically significant threshold) and *γ* (regularization parameter) are omitted from now on.

The idea behind the index *θ* is to gather the information about the *p-values* for different sample sizes against the predefined significance level *α*, usually equal to 5%. A distance *δ* (Eq. 9 in the Materials and Methods) is defined to compare the function *p*(*n*) with *α* for *all n* values. The distance *δ* measures the difference between the areas under the constant function at level *α* and the area under the curve *p*(*n*) (Fig. 1c). The distance *δ* is then used to obtain the binary index *θ* that indicates whether *p*(*n*) and the *α* constant are far from each other or not. If for most values of *n* the function *p*(*n*) is smaller than *α*, then *θ* = 1, which means that there are clinically reliable differences among the datasets being tested. Otherwise, *θ* = 0, which is interpreted as the non-rejection of the null hypothesis, and thus, the compared experimental set ups behave in a similar way.

As the exponential function is defined for all values *n* ∈ (−∞, +∞), it is necessary to determine a range of ri for which the function *p*(*n*) is meaningful in the context of this study. The decay of *p*(*n*) is concentrated in a range between *n* = 0 and a certain value of ri for which *p*(*n*) ≈ 0 (convergence of *p*(*n*)); so, *δ* should be only calculated in that range. A parameter *γ* is used as a regularizer to measure the sample size of convergence *n* = *n_γ_*, such that *p*(*n* = *n_γ_*) ≈ 0 (Fig. 1c and Eq. 8 in the Materials and Methods). Small *γ* values imply less restrictive decisions, i.e. *θ* = 1 when the groups being compared do not show clear differences. Nonetheless, the experimental evaluation of the method over synthetic and real data evidences *γ* = 5*e*^−06^ to be a reasonable choice (detailed information is given in the Materials and Methods and the Supplementary Material). Note that when *p*(*n*) is determined simply by the definition of the parameters *a* and *c* in Eq. 1, the minimum sample size needed to observe statistically significant differences at *α*-level can also be provided. As *p*(*n*) continuously decreases, the value of *n* for which *p*(*n*) is always smaller than *α* can be calculated easily. This value is called *n_α_* (Fig. 1c and Eq. 12 in the Materials and Methods).

## RESULTS

The decision index *θ*, descriptive parameters *a* and *c* (Eq. 1) and minimum data size *n_α_* provide an intuition about the distance between the distributions of the datasets being compared. To illustrate this, sample data generated from different normal distributions were compared using the Mann-Whitney U statistical test (Mann and Whitney, 1947) assuming a significance level *α* of 0.05 (Table 1 in Materials and Methods). When *N*(0, 1) is compared with *N*(0, 1), *N*(0.01, 1) and *N*(0.1, 1), *θ* is null; so those distributions are assumed to be equal if our approach is used. In the remaining comparisons though, according to our approach, *θ* = 1, thus there exist differences between *N*(0, 1) and *N*(*μ*, 1) for *μ* ∈ [0.25,3] (Fig. 1d). Together *a* and *c* provide a spatial representation of the distance between all the normal distributions and *N*(0, 1) (Fig. 1b and 1d). Likewise, when *N*(0, 1) is compared with *N*(*μ*, 1) for *μ* ∈ [0.1, 3], the value of *n_α_* increases as the mean value *μ* decreases. Indeed, *n_α_* cannot be determined when *N*(0, 1) is compared with *N*(0, 1) and *N*(0.01, 1), as the null hypothesis in this case is true. Therefore, *p*(*n*) is a constant function, which represents the uniform distribution of *p-values* under the null hypothesis (Figs. 1e and 1f, and Fig. 3 in Materials and Methods).

**Table 1.**
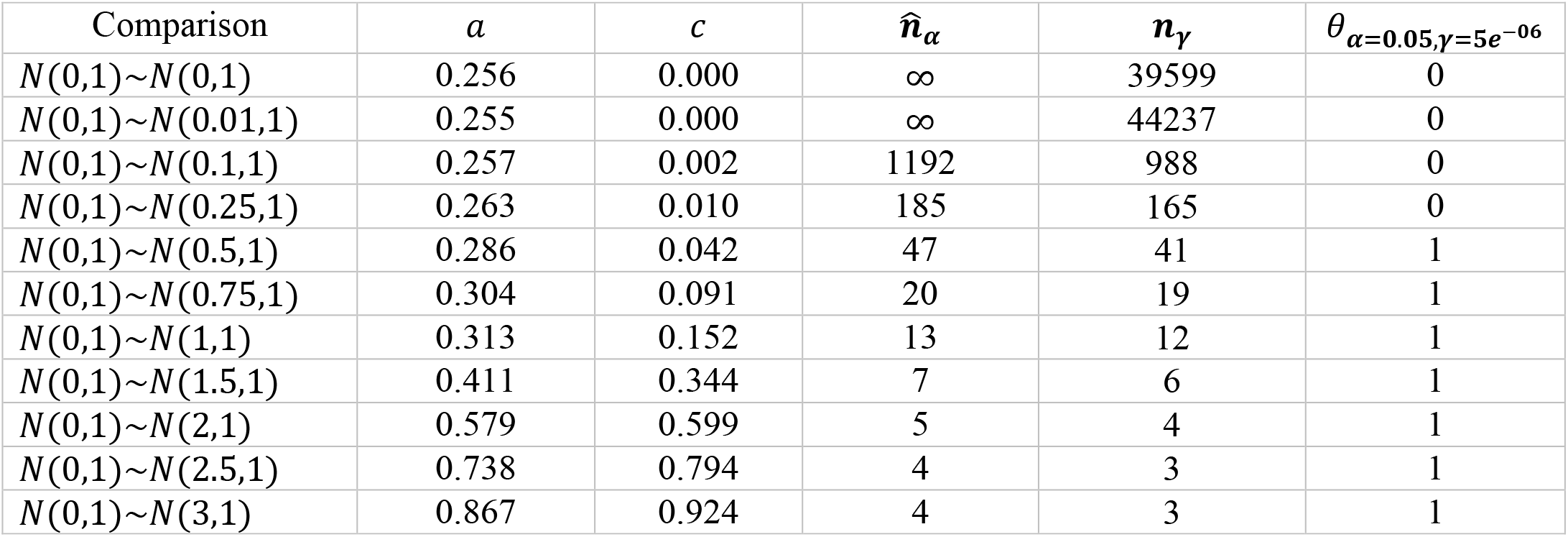
Parameters of the function p(n) after the exponential fit with α = 0.05 and γ = 5e ^−06^, for the comparison of a normal distribution with mean value 0 and standard deviation 1, and normal distributions of mean value 0, 0.01, 0.1, 0.25, 0.5, 0.75, 1, 1.5, 2, 2.5 and 3.

| Comparison | $a$ | $c$ | $\hat{n}_\alpha$ | $n_\gamma$ | $\theta_{\alpha=0.05, \gamma=5e^{-06}}$ |
| --- | --- | --- | --- | --- | --- |
| $N(0,1) \sim N(0,1)$ | 0.256 | 0.000 | ∞ | 39599 | 0 |
| $N(0,1) \sim N(0.01,1)$ | 0.255 | 0.000 | ∞ | 44237 | 0 |
| $N(0,1) \sim N(0.1,1)$ | 0.257 | 0.002 | 1192 | 988 | 0 |
| $N(0,1) \sim N(0.25,1)$ | 0.263 | 0.010 | 185 | 165 | 0 |
| $N(0,1) \sim N(0.5,1)$ | 0.286 | 0.042 | 47 | 41 | 1 |
| $N(0,1) \sim N(0.75,1)$ | 0.304 | 0.091 | 20 | 19 | 1 |
| $N(0,1) \sim N(1,1)$ | 0.313 | 0.152 | 13 | 12 | 1 |
| $N(0,1) \sim N(1.5,1)$ | 0.411 | 0.344 | 7 | 6 | 1 |
| $N(0,1) \sim N(2,1)$ | 0.579 | 0.599 | 5 | 4 | 1 |
| $N(0,1) \sim N(2.5,1)$ | 0.738 | 0.794 | 4 | 3 | 1 |
| $N(0,1) \sim N(3,1)$ | 0.867 | 0.924 | 4 | 3 | 1 |

**Figure 3.**
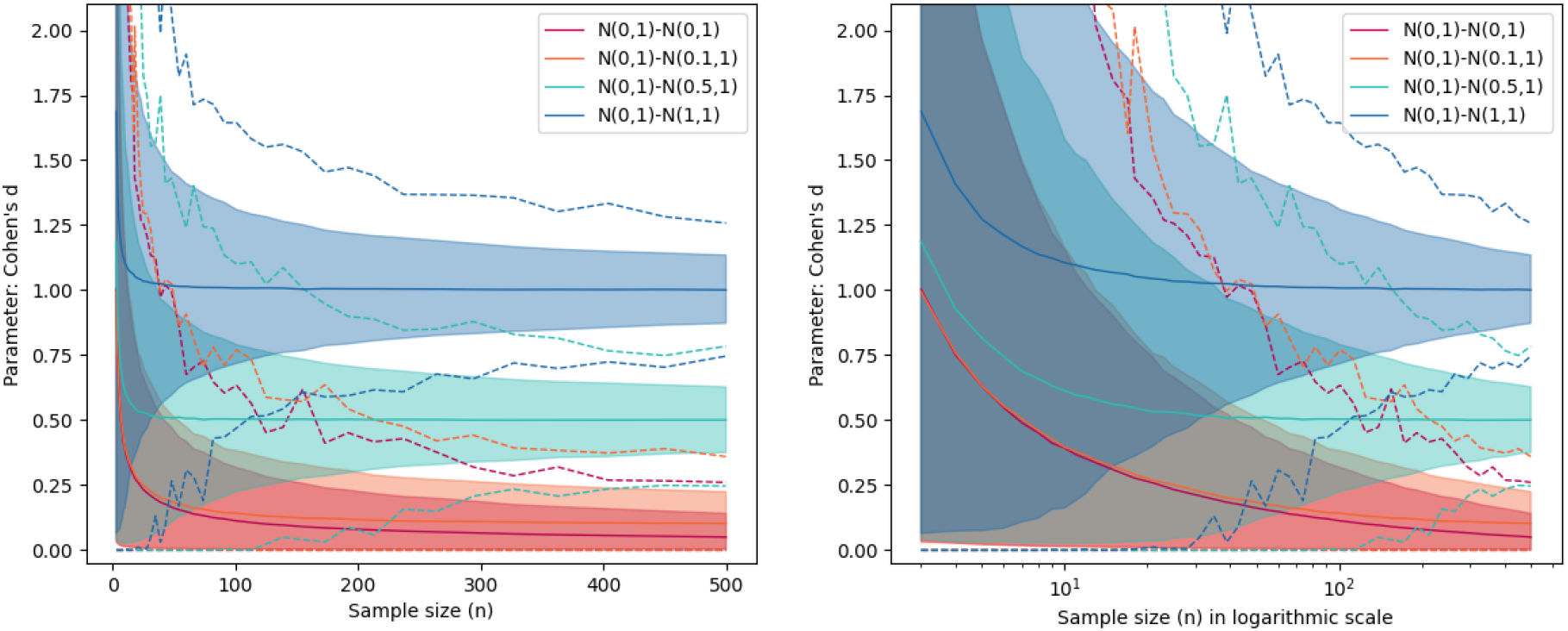
Estimation of the Cohen’s d value between a normal distribution with standard deviation of 1 and mean value 0, and normal distributions with standard deviation of 1 and mean values of 0, 0.1, 0.5 or 1. For each value of the sample size, we compute the Cohen’s d between simulated datasets from the two compared normal distributions 15000 times. The final confidence interval is obtained by clipping 95% of the values among the 15000 (filled area). The dashed lines show the maximum and minimum Cohen’s d values obtained for each sample size. The information is shown both in linear and logarithmic scale.

To prove the benefit of the proposed method in real data, we tested its different functionalities on published and non-published data from biological experiments. The first application of the method consists of discriminating between experimental conditions. In this case, we wanted to determine whether cancer cells cultured in 3D collagen matrices and imaged under a light microscope changed shape after administration of a chemotherapeutic drug (Taxol) (details about data collection and processing are given in the Supplementary Material). This information is relevant as it could give an indication about the metastatic potential of the treated cells (Wu et al., 2016, 2020). Three different groups were compared: control cells (non-treated), and cells treated with 1 nM and 50 nM Taxol respectively. Cells exposed to low concentrations of Taxol (1 nM) remained elongated (low roundness index, which suggests higher metastatic behavior of the cells), i.e. *θ* = 0 for the comparison between control cells and those treated with Taxol at 1 nM. However, when the dose was increased to 50 nM Taxol, cells became circular (lower metastatic behavior); therefore *θ* = 1 when comparing cells treated with 50 nM Taxol versus control cells, or cells treated with 1 nM Taxol (Fig. 2a and Table S3 in the Supplementary Material).

Secondly, we analyzed the flow cytometry data used by Khoury *et al*. (Khoury et al., 2018) to determine the transcriptional changes induced by the in vivo exposure of human eosinophils to glucocorticoids. The eosinophils belong to 6 different healthy human subjects. The proposed method allowed us to discriminate between treated and untreated eosinophils using the entire dataset. For that, we analyzed the eosinophil surface expression of the gene CXCR4 2 h after the exposure to 20 and 200 mcg/dL of Methylprednisolone. With the estimation of the function *p*(*n*) (Eq. 1), it is possible to conclude that the exposure of eosinophils to glucocorticoids causes a differential expression of CXCR4 (Fig. 2b), i.e. *θ* = 1 for the comparison between vehicle and eosinophils treated with 20 and 200 mcg/dL (Table S6 in the Supplementary Material). Indeed the conclusion is the same as the one made in (Khoury et al., 2018), where only the median fluorescence intensity of the data from each subject was calculated and the resulting 6 data points were compared (Fig. 2b). However, the latter approach can lead to false conclusions when the data distribution differs or when the data deviation is large.

The last use of the method we propose here consists of analyzing whether a single specific feature of the data (variable) can fully characterize the problem at hand. Many different biomolecular and biophysical features of human cells were analyzed in (Phillip et al., 2017) with the aim of predicting cellular age in healthy humans. The cells used in Phillip et al. study were collected from human subjects from 2 to 96 years-old. The method proposed in this manuscript can help to decide which features contain relevant information about subject aging. To show that, we re-analyzed the information of nuclei morphology and cell motility collected by Philip *et al*. (Phillip et al., 2017). The former is a large dataset and the latter is a small one. The information of 2-year-old human cells (the youngest one) was compared with the rest of the ages. The decay of *p*(*n*) in cell nuclei area and short axis length show that these nuclei morphology parameters are directly related to the age of human cells. The parameter *c* (Eq. 1) of the orientation of the cell nuclei is null in all cases, which indicates that this measure does not contain information about aging (Fig. 2c and Table S5 in the Supplementary Material). It is relevant to observe that the pattern in the plots of *a* and *c* indicates whether the analyzed feature can characterize the age of the patients: increasing values of *a* and *c* as the age of the patients increases (cell nuclei area and short axis length) and a null *c* regardless the age of the patient (orientation of the cell nuclei) (Fig. 2c). The estimated function *p*(*n*) for the total diffusivity of the cells of 2 year-old and 3 year-old human donors shows that even if a larger dataset was given, the result will remain the same (Fig. 2d and Table S4 in the Supplementary Material). Namely, *p*(*n*) does not decrease, therefore, there is strong evidence that the null hypothesis is true (i.e. *θ* = 0, groups behave similarly). So, in this case, the analysis of a small number of cells is enough to conclude the non-rejection of the null hypothesis. The most extreme cases given by the differences between 2 and 96 year-old human donors, can also be detected without the need of large datasets, *n_α_* = 11 (Fig. 2d). That is, the estimation of *p*(*n*) supports the decision about how many experimental samples need to be collected to conclude about the biological or clinical relevance of the differences between experimental groups.

The use of MCCV with large enough datasets guarantees robust estimators. In this case, different combinations of model parameters (MCCV iterations, used sample sizes and *γ* value) where repeatedly tested to evaluate the variability of the decision index (*θ*) and its sensitivity to the method set up (Test of reliability in the Materials and Methods). A larger *γ* value results in a more restrictive decision index *θ* in the task of detecting interesting differences (Table S7 and S9 in the Supplementary Material). When the number of iterations of MCCV is drastically reduced, the decision index (*θ*) shows instability only in those cases for which it is not clear that the groups differ from each other (Table S8 and S10 in the Supplementary Material)

## DISCUSSION

The use of statistical hypothesis testing is largely extended and well established in the scientific research literature. Moreover, the number of statistically significant *p-values* reported in scientific publications has increased over the years (Chavalarias et al., 2016) and there exists a tendency among researchers to look for that combination of data that provides a *p-value* smaller than 0.05 (Bruns and Ioannidis, 2016). However, it has been shown here and also by Altman and Krzywinski, 2017; Amrhein et al., 2019; Bruns and Ioannidis, 2016; Halsey et al., 2015; Nuzzo, 2014, that the assessment of the *p-value* has some drawbacks which can lead to spurious scientific conclusions. The data recorded from high-content, high-throughput studies, and the capacity of the computers to analyze thousands of numbers, has enabled us to enlighten the current uncertainty around the exploited *p-value*.

We report clear evidence about the well-known dependence of the *p-value* on the size of the data (Altman and Krzywinski, 2017; Krawczyk, 2015; Lin et al., 2013). This particular feature of the *p-value* is used to characterize the differences among the groups of datasets being analyzed. Due to the lack of techniques that exploit the sensitivity of the *p-value* with respect to the sample size, we believe that our method will have a huge impact in the way scientists perform hypothesis testing.

With the proposed estimation of the decay of the *p-value* with the sample size, we provide a new perspective about hypothesis testing that prevents from treating the *p-value* as a dichotomous index. Using a simple mathematical formulation, an unbiased decision index *θ* is defined to enable good *praxis* in the same context as statistical hypothesis testing. The method takes advantage of large sample sizes to analyze the dependence of the *p-value* using cross validation. This approach provides stable measures that are robust to the noise in the data or the uncertainty around the decision making process. Indeed, the presented method is applicable in any field of study beyond life sciences as the classical NHST. Moreover, this methodology could be transferred to multiple comparison frameworks such as the ANOVA test by approximating the *p-value* function for each pair of comparisons. The proposed approach used as a preliminary analysis, provides evidence about the existence (or not) of real differences from a practical perspective, even when large datasets are not available. Therefore, it supports the management of new data collection and can help researchers to reduce the cost of collecting experimental data.

The decision-index *θ* obtained with the proposed analytic pipeline relies on a new threshold called *γ*, as shown in Materials and Methods. Compared to the classical *p-value* and *α* threshold, the parameter *γ* is mathematically constrained and *θ* is stable to its variations. Similarly, *n_α_* is an unbiased effect size indicator, i.e. how different the samples are or how big this difference is (further details about robustness are given in Materials and Methods). Additionally, the fitted parameters *a* and *c* that determine *p*(*n*) in Eq. 1, represent graphically how each of the conditions of an experiment relate to each other regarding the distribution of their values (Fig 1b, 1d and 2a-c). When the differences between the compared samples increase, the value of *a* and *c* increase as well regardless of the sample size, which suggests again, that *a* and *c* are unbiased indicators of the effect size.

The computational cost of the proposed data diagnosis increases proportionally with the number of groups to compare and the numerical setup of MCCV. Therefore, the optimization of the code and its connection to either a GPU or cloud computing is recommended. Overall, we advocate for the implementation of our pipeline in user-friendly interfaces connected to either cloud-computing or GPU. The code provided within this manuscript is built into the free software Python, so that anyone with limited programming skills can include any change to obtain a customized tool.

## ACKNOWLEDGEMENTS

This work was produced with the support of the Spanish Ministry of Economy and Competitiveness (TEC2015-73064-EXP, TEC2016–78052, PID2019-109820RB-I00) (AMB), a 2017 Leonardo Grant for Researchers and Cultural Creators, BBVA Foundation (AMB), and grants from the US National Institutes of Health (UO1AG060903 (DW, JMP), P30AG021334 (JMP) and U54CA143868 (DW)). We also want to acknowledge the support of NVIDIA Corporation with the donation of the Titan X (Pascal) GPU used for this research. This material is based upon work supported by the National Science Foundation Graduate Research Fellowship under Grant No. 1746891. We thank Claire Jordan Brooks, Prof. Joachim Goedhart (University of Amsterdam), Laura Nicolás-Sáenz, and Pedro Macías-Gordaliza for fruitful discussions. We also want to acknowledge the valuable comments from the reviewers of this manuscript.

## AUTHOR CONTRIBUTIONS

E.G.M. contributed to the conception and the implementation of the mathematical method, designed the data analysis and wrote the manuscript with input from V.G., D.W. and A.M.B.. A.S., H.J., J.M.P. and D.W. contributed with the experimental data. All authors contributed to the interpretation of the results. All authors reviewed the manuscript.

## MATERIALS AND METHODS

### Available in the following pages

- DATA DETAILS
  - Drug analysis on phase contrast microscopy
- METHOD DETAILS
  - *p-value* as an exponential function of data size
  - Distance to the *α*-level of statistical significance
  - Mathematical formulation of the decision index
    - Restricting an optimal threshold
  - Data characterization in stable and uncertain cases.
  - Test of reliability
- IMPLEMENTATION DETAILS

## SUPPLEMENTAL INFORMATION

available in the following pages.

## CODE AVAILABILITY

at https://github.com/BIIG-UC3M/pMoSS.

## Materials and Methods

### DATA DETAILS

We describe the non-published dataset that corresponds to the first example in the main text.

#### Drug analysis on phase contrast microscopy

Phase contrast microscopy images of a human invasive ductal carcinoma (MDA-MB-231) cell line were acquired. The set-up used was composed by a Cascade 1K CCD camera (Roper Scientific), mounted on a Nikon TE2000 microscope with a 10X objective lens. Cells were embedded in 3D collagen type I matrix at 100.000 cells/mL. The time lapse videos were recorded every two minutes with a focus plane of at least 500 *μm* away from the bottom of the culture plates to diminish edge effects (He et al., 2017). Three different groups of cells were analyzed: control and treated with fresh media at 1 nM Taxol and 50 nM Taxol. Ten videos of 16.5 hours (500 frames of 809 *μm* x 810 *μm* with a resolution of 0.806 *μm/pixel*) each were analyzed per group.

All videos were automatically processed using a convolutional neural network (U-net (Ronneberger et al., 2015)) to get binary masks for the cell bodies and their protrusions. The resulting semantic segmentation corresponds uniquely to focused cells in the image. For each of these cells, their body and protrusions are segmented. See some examples of the resulting segmentation in Figure S2.

Using the segmentations, eight different morphological measurements were calculated: cell body size (CS), cell body perimeter (CP), cell body roundness (CR), cell with at least one protrusion (Pb), protrusion size (PS), protrusion perimeter (PP), protrusion length (PL) and protrusion diameter (PD) (Table S1). Further information about the distribution of each of the measurements is given in the Supplementary Material. Same as the analysis done for CR (Fig. 2a in the main manuscript), the differences between control and 1 mM and 50 nM Taxol were analyzed using each of the remaining variables (Figures S4-S6 and Table S3).

### METHOD DETAILS

Here, we first show the relation between the accuracy of an estimator and the size of the data being analyzed. Then, we provide the mathematical details behind our hypothesis that the *p-value* is a random variable that critically depends on the size of the sample and that the *p-value* function can be approximated with an exponential function of the sample size *n*. With this idea in mind, we define the method of how to work with the *p-value* as a function and to determine when a statement of practical significance can be made (*θ_α,γ_*, Eq. 9). Once the problem is described technically, it is possible to calculate the minimum size *n_α_* at which the null hypothesis of the test is statistically significant (Eq. 11). This parameter *n_α_* can be used to characterize the data. Finally, the reliability of our method is rigorously tested.

#### The effect of the data size in empirical estimators

Fig. 1–2 illustrate that the precision on the mean estimators increase with the size of the data (Krzywinski and Altman, 2013c; Lin et al., 2013). For each sample size, we use Monte Carlo with simulated samples from normal distributions to calculate both the mean and its two sided 95% confidence interval (CI). Note that we distinguish between the theoretical CI (Fig. 1) and the bootstrapped CI (Fig. 2). Following the same procedure, two of the most common measures of effect sizes are also calculated: Cohen’s d and Jensen-Shannon divergence and distance, Fig 3–4 respectively. We compute both, their mean value for all the computations at each fixed sample size and the bootstrapped two-sided 95% CI. Fig.1–4 do also display the maximum and minimum simulated values for each sample size. In all cases, there is an exponential convergence to the estimated value.

**Figure 4.**
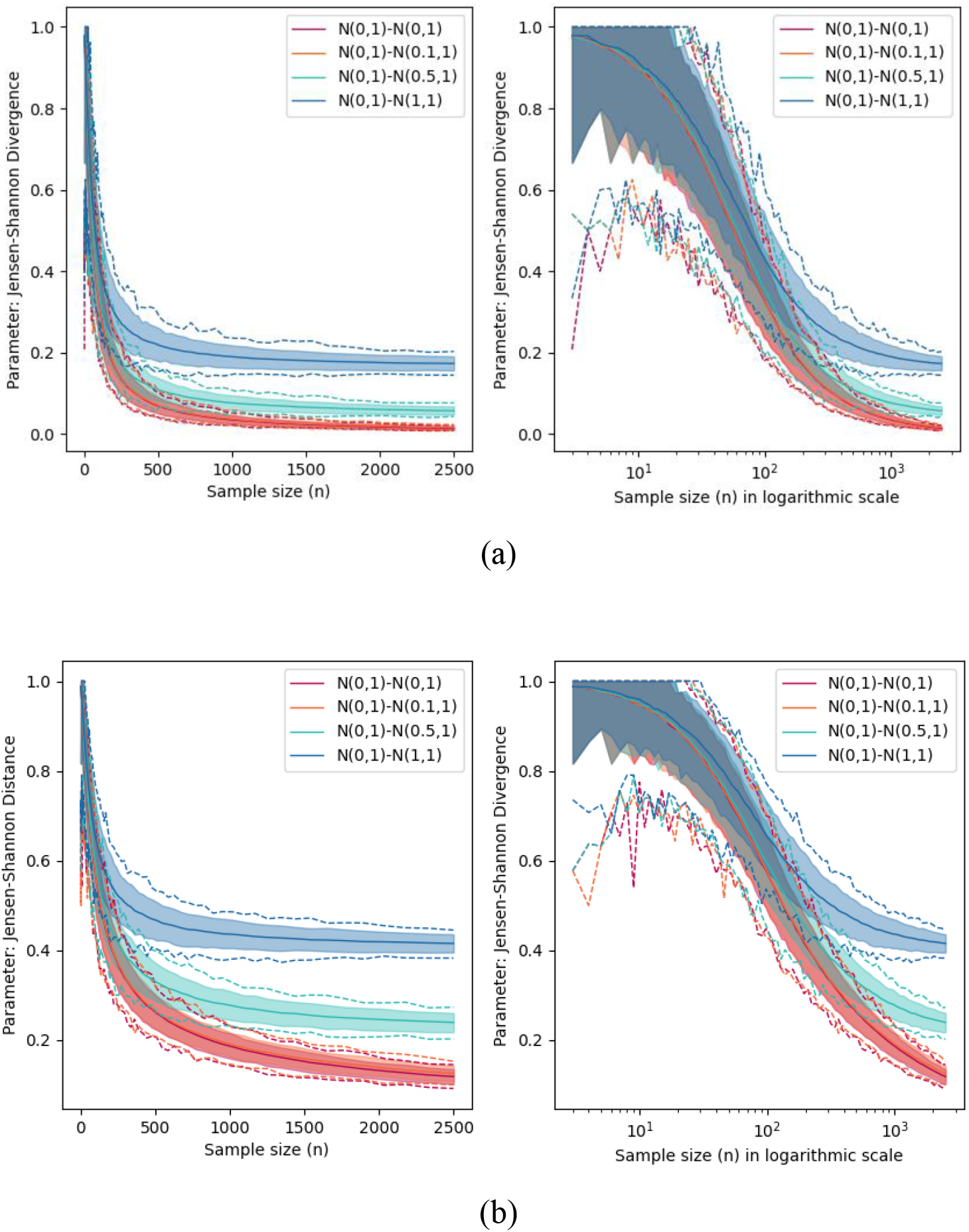
Estimation of the (a) Jensen-Shannon divergence and (b) distance values between a normal distribution with standard deviation of 1 and mean value 0, and normal distributions with standard deviation of 1 and mean values of 0, 0.1, 0.5 or 1. For each value of the sample size, we compute the Jensen-Shannon divergence and distance between simulated datasets from the two compared normal distributions 15000 times. The final confidence interval is obtained by clipping 95% of the values among the 15000 (filled area). The dashed lines show the maximum and minimum Jensen-Shannon divergence and distance values obtained for each sample size. The information is shown both in linear and logarithmic scale.

#### *p-value* as an exponential function of data size

Fig. 5 illustrates the idea that the *p-value* is a function that depends on the sample size π. There exists a continuous inverse relation between *p-values* and π, i.e. *p*-values decrease when *n* increases, (Altman and Krzywinski, 2017; Krzywinski and Altman, 2013a, 2013b). This allows us to assume that *p-values* can be considered indeed, as a function of *n*, i.e. *p*(*n*).

**Figure 5.**
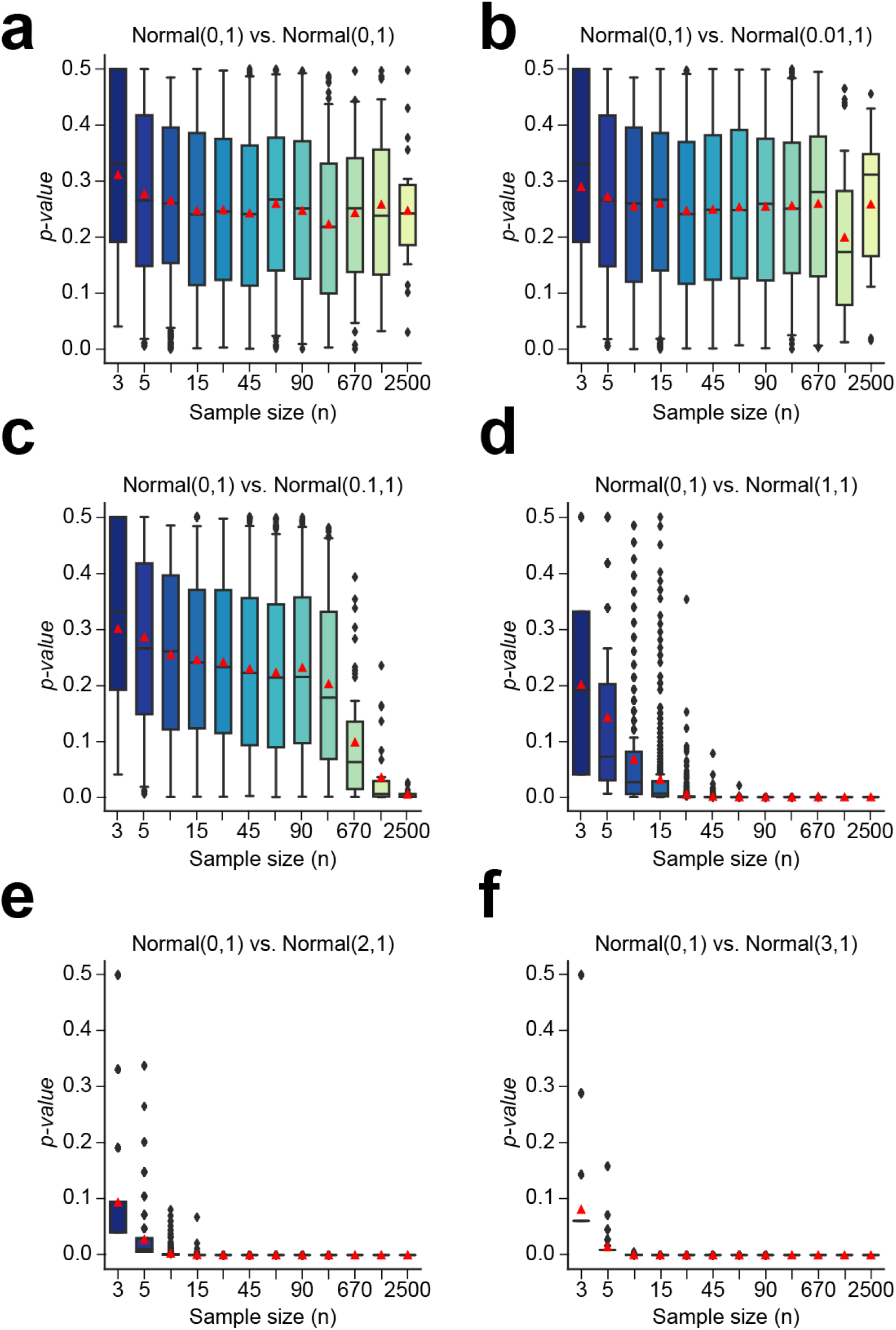
Distribution of the *p-values* obtained when two normal distributions are compared. For each fixed value of the sample size (3, 5, 10, 15, 30, 45, 60, 90, 200, 670, 1750 and 2499 points), two normal distributions of that size are simulated and compared via the Mann-Whitney statistical test. This procedure is repeated multiple times. A normal distribution with a mean of 0 and a standard deviation of 1 is compared with a normal distribution of mean: **(a)** 0, **(b)** 0.01, **(c)** 0.1, **(d)** 1, **(e)** 2, and **(f)** 3 and a standard deviation of 1. When both normal distributions are almost the same, **(a)** and **(b)**, the *p-value* follows a uniform distribution. Though, as long as both normal distributions get farther to each other, the distribution of *p-values* become closer to a normal distribution with a faster decay.

Either with Mann Whitney U test (Mann and Whitney, 1947) or with Student’s *t*-test (Student, 1908), it can be proved that the obtained *p-value* converges to zero when the sample size is large and the distributions being assessed are not exactly the same, i.e., the *p-value* tends to zero when the sample size tends to infinity. A mathematical demonstration of this statement is available in the Supplementary Material.

Going a step further, we claim that the *p-values* can be indeed written directly as a function of *n*, *p*(*n*), and that this function adjusts well to an exponential function. To show this, we first estimate the value that the *p-value* function has at each possible value of *n*. This can be done easily with the Monte Carlo cross validation method (MCCV) (Xu and Liang, 2001): at each iteration *i* of the procedure, *n* = *n_i_* is fixed, and two populations of size *n_i_* are compared. This procedure is repeated many times in each given iteration *i* to cover the variability of the problem at *n* = *n_i_*. At the end, we have as many sets of *p-values* as iterations *I* that are of the form:

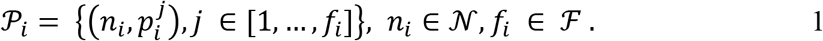

where 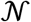 and 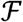 are grids of natural numbers defined according to the sample size, so the computational cost of MCCV is reduced without losing information. Further details are given in the description of the MCCV routine in the Supplementary Material. Note that this procedure is similar to the upstrap (Crainiceanu, 2018) using an increasing fraction of the sample. The details about the procedure followed for the estimation of the *p-values* are explained in the Supplementary Material.

In Fig. 5, the procedure is applied using random populations from different normal distributions. We distinguish two different situations: either the obtained distributions are uniform, so the mean value of all the 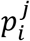 values is constant for any *i* (Figs. 5a and 5b); or the mean value tends to decrease when the sample size *n* increases (Figs. 5c-f). In other words, *p*(*n*) can be written as a continuous function. Hence, for each iteration *i*, each set of *p_i_* values is averaged to obtain the empirical estimation of the function *p*(*n*) at *n* = *n_i_* (red markers in Fig. 5). Then, a smooth curve is fitted to these values using locally weighted scatter plot smoothing (LOWESS) (Cleveland, 1979), which shows *p*(*n*) has an exponential shape (Figs. 6a and 6b).

**Figure 6.**
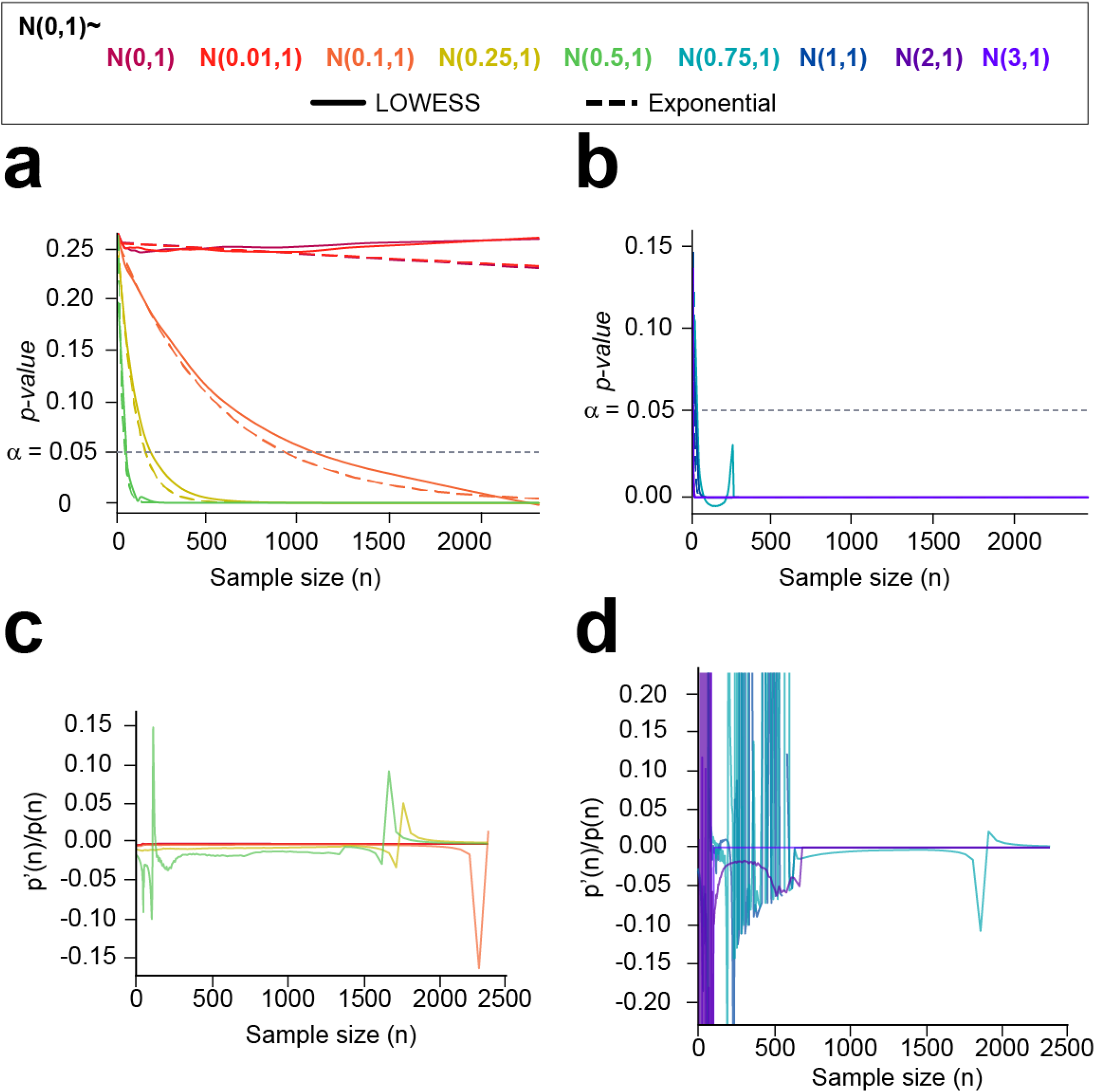
A normal distribution with a mean of 0 and a deviation of 1 is compared with a normal distribution of mean (0, 0.01, 0.1, 0.25, 0.5, 0.75, 1, 2 and 3). Multiple *p-values* are calculated for a sample size between two and 2500 (Fig. 3). **(a)** and **(b)** Locally weighted scatter plot smoothing (LOWESS) fit to the mean *p-values* (red markers in Fig. 3) computed for each value of the sample size *n*. Likewise, an exponential function is fitted to all the simulated *p-values*. **(c)** and **(d)** Quotient between each LOWESS curve and its differential. **(c)** Comparison of *N*(0, 1), with *N*(0, 1), *N*(0.01, 1), *N*(0.1, 1), *N*(0.25, 1) and *N*(0.5, 1). **(d)** *N*(0, 1) is compared with *N*(0.75, 1), *N*(1, 1), *N*(2, 1) and *N*(3, 1). Constant quotients and accurate exponential fits show empirically that *p*(*n*) has an exponential nature.

To prove that the estimated function *p*(*n*) can be written as an exponential function, it is sufficient to verify that the quotient between its first derivative *p*′(*n*) and the function *p*(*n*) is itself a constant, i.e.

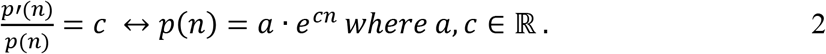

Collecting the values *p*(*n*) of the LOWESS fit, the quotient 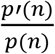 is calculated (Figs. 6c and 6d). Most of the quotients verify the condition in Eq. 2. In Fig 6c, we show cases in which it is more challenging to decide whether there exists a statistical difference, as for instance, when *N*(0, 1) and *N*(0.1, 1) are compared. When *p*(*n*) becomes very small, the quotient 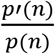 has more outliers, especially when the sample size *n* is small. This can be observed when comparing *N*(0, 1) with *N*(0.75, 1), *N*(1, 1), *N*(2, 1) and *N*(3, 1). (Fig. 6d). These are extreme cases in which there exist clear differences between populations and therefore, *p-values* are close to zero most of the time.

As we have proved above that the estimated function *p*(*n*) can be written as an exponential function, an exponential curve is fitted to all the pairs of values 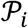 calculated with MCCV (Figs. 6a and 6b). Both LOWESS and exponential curves are very close to each other, even if the former was fitted using the mean values of each group 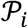 and the latter with all of them. An exponential fit is more suitable in this case as it is calculated with all the values obtained through MCCV, and only outputs positive values by definition. A LOWESS approximation can occasionally lead to biased negative values, such as when *N*(0, 1) and *N*(0.75, 1) are compared while the *p-values* are positively defined. Note that as *p*(*n*) → 0 when *n* → ∞, *c* < 0 necessarily in Eq. 2. Therefore, we assume from now on that *p*(*n*) can be given as an exponential function of the form

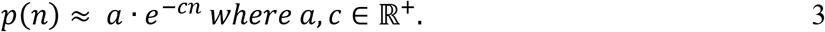

Here the parameters *a* and *c* control the amplitude and the decay of the function *p*(*n*), respectively. If *c* = 0, then the value of *p*(*n*) would be uniform in *a*: *p*(*n*) = *a*. As *p-values* are computed probabilities and the global maximum of *p*(*n*) is *a, a* belongs to the [0, 1] interval.

#### Distance to the *α*-level of statistical significance

The ideal case of a true (1 – *α*) statistical significance would lead to the rejection of the null hypothesis independently of data size, i.e., (1 – *α*)·100% of *p-values* would always be lower than *α*. Hence, we claim that whenever there exist clinically meaningful differences between two samples, *p*(*n*) reaches *α* rapidly. So, the values of *p*(*n*) are mostly distributed in a range smaller than *α*. Therefore, we compare all the values of the curve *p*(*n*) with *α*(*n*) = *α*. In the discrete case, we would evaluate *α* – *p*(*n* = *n_i_*) for each index *i* and sum all the results: if the sum is positive, then *p*(*n*) is smaller than *a* most of the time. In the continuous case, this sum is obtained by integrating the difference

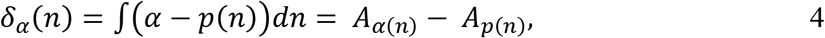

where *A_α_* is the area under the constant function *α* and *A*_*p*(*n*)_ is the area under the estimated *p-values’* curve, *p*(*n*) (Fig. 7). A positive *δ*(*n*) implies that *A_α_* is larger than *A*_*p*(*n*)_, i.e. most of the values in *p*(*n*) are below the significance threshold *α*; a negative *δ*(*n*) implies the opposite.

**Figure 7.**
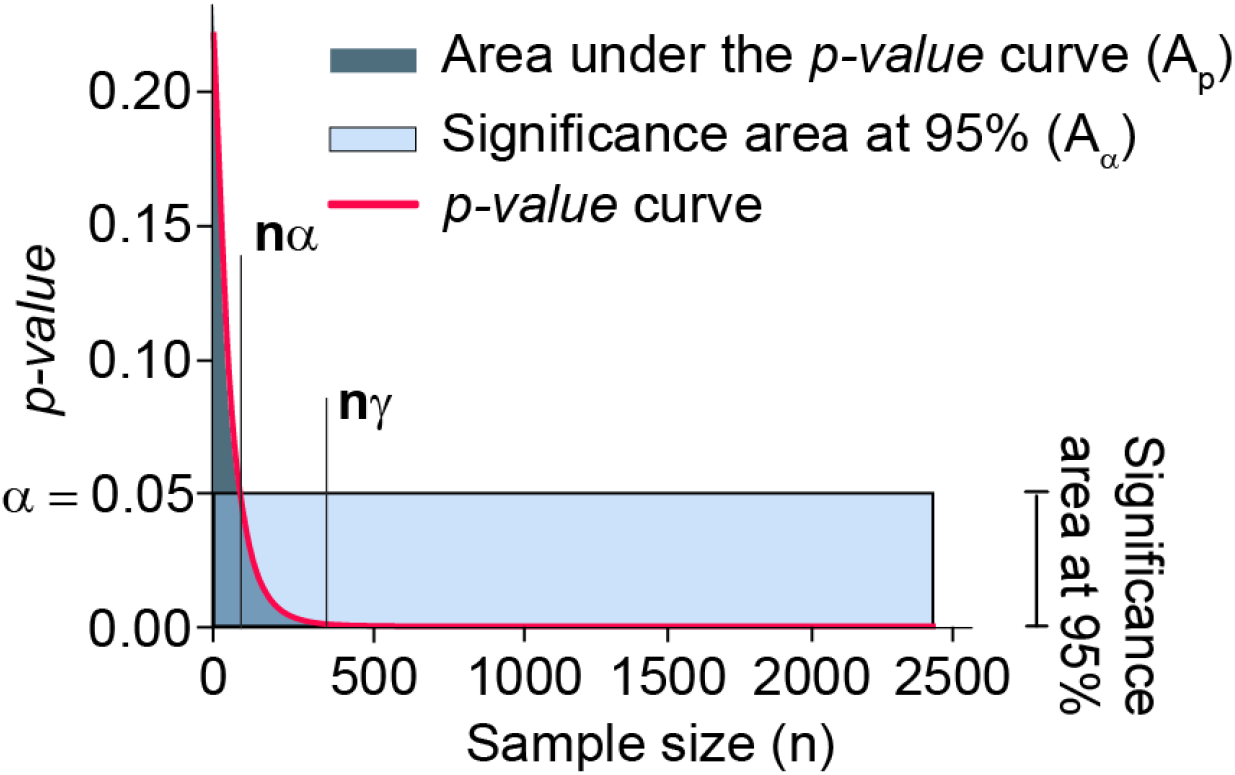
Comparison of a 95% of statistical significance (*α* = 0.05) and an *n*-dependent *p-value* curve. The parameter *n_α_* represents the minimum sample size to detect statistically significant differences among compared groups. The parameter *n_γ_* represents the convergence point of the *p-value* curve. When the *p-value* curve expresses statistically significant differences, the area under the red curve (*A*_*p*(*n*)_) is smaller than the area under the constant function *α* = 0.05 (*A*_*α*=0.05_) when it is evaluated between 0 and *n_γ_*.

As shown in the next paragraphs, Eq. 4 aims to quantify and evaluate the distribution of *p-values* (i.e., the distribution of {(*n*, *p*(*n*)), *n* ∈ *N*}) taking into account two aspects, whether (1) most of the *p-values* are smaller than *α* and (2) the decay of *p*(*n*) is large enough.

#### Mathematical formulation of the decision index

By means of the exponential expression of *p*(*n*) given in Eq. 3, the measure *δ_α_*(*n*) (Eq. 4) can be rewritten as follows

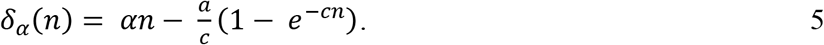

Due to the limits of *α* and *c*, *δ_α_*(*n*) is still well-defined. However, in the limit of *n*, *δ_α_*(*n*) will always be positive and it tends to infinity:

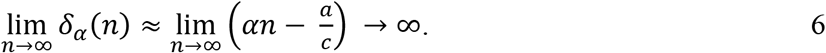

Also, from a practical perspective, the area of interest to evaluate the decay of *p*(*n*) is that enclosed between zero and its convergence point *n*: |*p*′(*n*)| ≈ 0. Namely, a relevant sub-sample of size *n* can be computed as

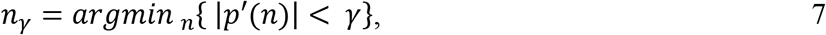

where *γ* is the threshold chosen to determine the convergence point (Fig. 7). Finally, *δ_α,γ_* is now formally defined as

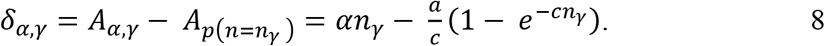

As claimed at the end of the last section, the computation of *δ*_*α,γ*_ enables the identification of a rapid convergence to zero at small values of *n* induced by the fast decay of *p*(*n*), which is indicative of the existence of relevant differences from a practical perspective.

The decision index we propose, *θ_α,γ_*, is defined as

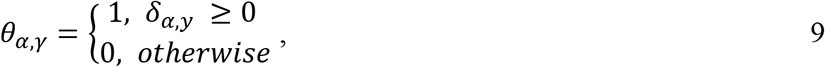

where *δ_α,γ_* follows Eq. 8.

#### Delimiting the convergence of the curve *p*(*n*)

The proposed approach depends on two thresholds: (1) significance threshold *α* and (2) the convergence threshold *γ*. The former measures the level of statistical significance, while the latter controls decisions. Therefore, the only critical threshold to discuss in this work is *γ*.

The rules to follow for the selection of the threshold *γ* are:

- The parameter *a* is the maximum value that *p*(*n*) can take. Therefore, if *a* is smaller than *α*, then *θ*_*α,γ*_ = 1 for any *γ* given.
- As *δ*_*α,γ*_(*n*) tends to infinity with *n*, the smaller the value of *γ* is set, the larger *n_γ_* will be and the chances of *θ*_*α,γ*_ = 1 will also increase.
- The values of *γ* should be small: *α* is considered a significant number and *p*(*n*) values are constantly compared with it. It seems reasonable to compare the slope of *p*(*n*) at the convergence point with a value smaller than *α*, which is usually smaller than 0.1.

Eq. 7 implies

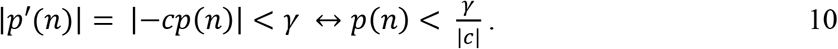

So, if *γ* is chosen such that 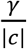 is greater than *α*, it would vanish the assumption that *p*(*n*) has arrived at a convergence point equivalent to zero. Therefore, our claim is that 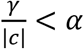 with at least *γ* < 0.1.

#### Data characterization in stable and uncertain cases

The threshold *γ* controls severe decisions and it is limited to 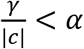 with at least *γ* < 0.1. In this section we study the range *γ* ∈ [1*e*^−12^, 0.1] to see the effect on the decision index *θ_αγ_*. Namely, the lower *γ* is set, the larger the value *n* is to determine *p*(*n*)’s convergence. Hence, when *γ* is small, the decision index will determine that there are practical differences among groups more often, becoming then less strict. In Fig. 8a, we show the dynamics of *θ*_*α*=0.05,*γ*_ when *γ* changes: the dark area (*θ*_*α*=0.05,*γ*_ = 1) increases inversely to *γ*, showing that the chances for which the null hypothesis is rejected increase as well. Moreover, the limit between dark and light (*θ*_*α*=0.05,*γ*_ = 0) areas is precisely the curve *δ_α,γ_* = 0. The value of *γ* determines this curve and therefore, the conditions for which *θ*_*α*,=0.05,*γ*_ = 1 (dark area) and *θ*_*α*=0.05,*γ*_ = 0 (light area). In Fig. 8b, we illustrate the condition *δ_α,γ_* = 0 when *α* = 0.05, as a function of *a, c* and *γ*. The case *γ* = 5*e*^−06^ is underlined in black.

**Figure 8.**
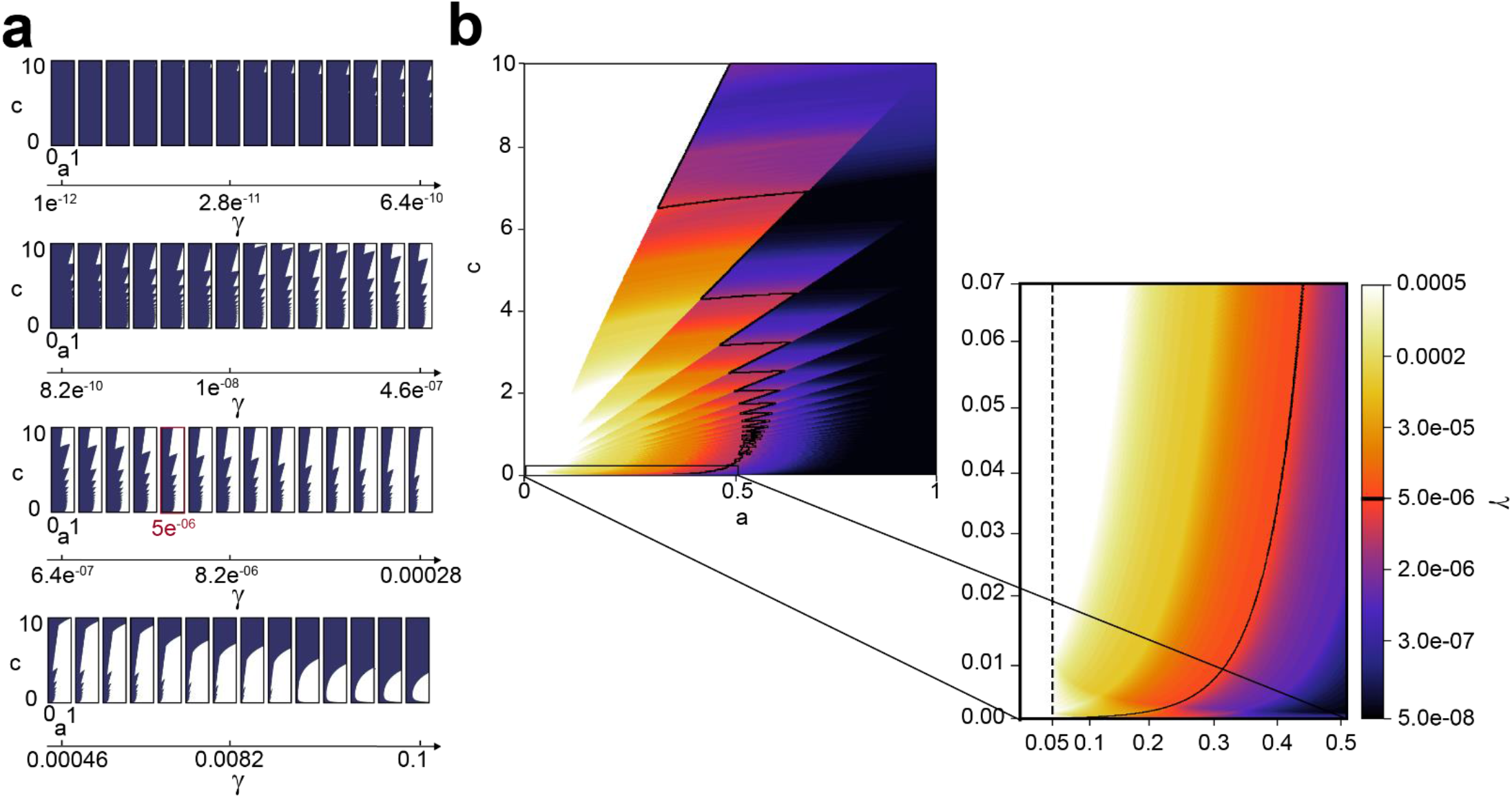
Decision index *θ*_*α*=0.05,*γ*_ for different values of parameters *a* and *c* in the function *ae^−cn^* and threshold *γ*: **(a)** Each of the subplots is drawn for a specific value of *γ*, being the dark area the cases for which we conclude that there are meaningful differences (*θ*_*α*=0.05,*γ*_ = 1), and white area the rest of the cases *θ*_*α*=0.05,*γ*_ = 0; **(b)** Colors in the image correspond to the values of *γ* for which *δ*_*α*=0.05,*γ*_ = 0. The black frontier shows *δ*_*α*=0.05,*γ*=5*e*^−06^_ = 0 (red box in **(a)**). All the values of *a* and *c* for which *θ*_*α*=0.05,*γ*=5*e*^−06^_ = 1 (practical differences) lie on the left side of this limit and, the rest, on the right. The plots shown in **(a)** show the influence of the parameter γ in a wide range of values, while the plots shown in **(b)** are limited to the range of values we find in this posterior experiment. The vertical dashed line indicates the cases *a* = 0.05 which are the cases in which *p*(*n*) outputs a 95% statistically significant value.

There exist some points (*a, c*) for which the rejection or not of the null hypothesis is independent of *γ*. A clear example is the case in which *a* ≥ *α* and *c* ≈ 0. These cases represent the situation in which the null hypothesis cannot be rejected with a statistical significance of level *α*. For instance, when *N*(0, 1) is compared with *N*(0, 1) or *N*(0.01, 1) (Fig. 1b in the main manuscript, Figs. 5a-b). Likewise, if *a* ≤ *α* or *c* is large enough, the null hypothesis is always rejected with a statistical significance of level *α*. For instance, when *N*(0, 1) is compared with *N*(2, 1) or *N*(3, 1) (Fig. 1b in the main manuscript, Figs. 5e-f).

The proposed methodology allows us to classify by their level of uncertainty the decisions on the differences among the groups of observations. Namely, if the differences can be considered relevant from a practical perspective or not. The parameters of the exponential curve in Eq. 1 in the main text determine the axis of any of the plots in Fig. 8. Therefore, once an exponential curve is fitted and parameters *a* and *c* are estimated, it is possible to know in which position of the graph the case of study is: clear cases will always be close to the left or to the right side of the graphs in Fig. 8, while most unstable or unclear cases will be placed in the middle. Therefore, with this method, it is possible to determine if there are clinically significant differences or not. When these differences are not sufficiently clear, it might be necessary to perform a deeper study.

An intuitive interpretation of statistically significant differences between two groups (the classical threshold *p-value* < *α*) is that their mean CI do not overlap. These CI decrease when the size of the data increases (Krzywinski and Altman, 2013c), Fig. 1–2. The rest of the section is devoted to study how large two populations must be in order to obtain non-overlapping CI. Interestingly, the estimation of the function *p*(*n*) allows us to determine the specific minimum value of *n*, *n_*α*_*, for which *p*(*n*) is lower than the significance level *α* (Fig. 8). This value is the solution to the equation

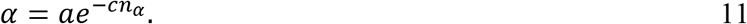

As computed, *n*_*α*_ represents the minimum sample size needed to obtain a statistically significant *p-value*, in case it exists. In other words, reproducing an experiment with *n_*α*_* samples assures the rejection of the null hypothesis. The estimated *n_*α*_* allows to assess the strength of the evidence against the null hypothesis. If *n_*α*_* is small, the strength of the statistical difference is very clear and two populations are distinguishable.

The parameters *a* and *c* in Eq. 11 are obtained empirically through MCCV so they can introduce some bias in the calculation of *n_*α*_*. Hence, a better estimator of *n_*α*_*, 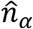, can be computed using the *p-values* obtained directly from the data and their variance

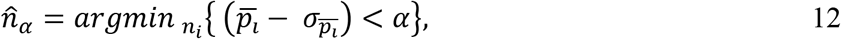

where 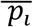 represents the mean of the set of values *p_i_* (MCCV) and 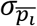, the mean standard error (SEM), which is included to correct for the variability of the estimated *p-values*. The estimator 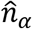 is limited to those cases in which the data is large enough: if the size of the data is smaller than *n_*α*_*, then 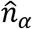 cannot be computed (Fig. 1e-f, Fig. 2d in the main manuscript). As 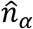 is more restrictive than *n_*α*_*, its value will always be slightly larger (Table 1). The values in Table 1 are similar to those values in Fig 1–2 for which the CIs do not overlap.

#### Test of reliability

Unlike many computational methods, the analysis of statistical significance of the differences between two groups cannot be evaluated by means of Ground Truth data, simulations or human-made annotations. Nonetheless, it is possible to determine the robustness on the reproducibility of the results. Namely, whether the decision taken about the stated null-hypothesis (*θ*_*α,γ*_) is maintained when the experiment is repeated. To do so, we test our method using simulated normal distributions.

Any data diagnosis carried out with the proposed method depends on the value *γ* chosen and the limitations posed by its computational intensive nature. As done at the beginning of this work, we compare the normal distribution *N*(0, 1) with *N*(0.01, 1), *N*(0.1, 1), *N*(0.25, 1), *N*(0.5, 1), *N*(0.75, 1), *N*(1, 1), *N*(2, 1) and *N*(3, 1). We should obtain *θ_α,γ_* = 1 when comparing the most similar distributions such as *N*(0, 1) and *N*(0.01, 1). In contrast, we should get *θ_α,γ_* = 0 when comparing the most different distributions, such as *N*(0, 1) and *N*(2, 1).

To evaluate the effect of *γ*, *p*(*n*) is simulated for all pairs of normal distributions and it is compared with a significance level of *α* = 0.05 using different values of *γ* (Table S7 in the Supplementary Material). The lower the convergence criterion *γ* is, the less restrictive the diagnosis is (Fig. 8). Using the simulated data, the range of *θ*_*α*,=0.05,*γ*_ values obtained let us recommend a value for this parameter. Note that *γ* satisfies 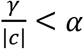 and that *c*, even if defined as a positive value, in our experiments is shown to be in the [0, 1] range (Table 1). When *N*(0, 1) and *N*(0.1, 1) are compared with a small *γ*, (*γ* = 2.5*e*^−06^), the decision index *θ*_*α*=0.05,*γ* = 2.5*e*^−06^_ = 1 indicates that there exist meaningful differences among both distributions, which is the opposite of what we expected. If the value of parameter *γ* increases, the decision index will output that the two compared groups do not display meaningful differences in those cases in which there is a larger uncertainty about this decision. For instance, when *N*(0, 1) and *N*(0.25, 1) are compared with *γ* = 5*e*^−05^, *θ*_*α*=0.05,*γ*=5*e*^−05^_ = 0. However, the latter is not straightforward for two reasons: *δ*_*α*=0.05,*γ* = 5*e*^−05^_ = –5.84 (small difference) and 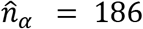 (few samples to observe statistically significant differences). As it is shown in Fig. 8, a value *γ* > 5*e*^−04^ results in *θ*_*α*=0.05*γ*_ = 0 for all the cases in Table 1. Note that the values of the function *p*(*n*) are enclosed in the [0, 1] range and that *γ* is used to determine where the *elbow* of this function *p*(*n*) lies, i.e. the convergence point. Therefore it is reasonable to use the same value of *γ* regardless the data that is being analyzed. With the results of the evaluation, we strongly recommend the use of *γ* = 5*e*^−06^. Indeed the decisions about having interesting differences among the compared groups are robust to the changes in the value of *γ* in both simulated and experimental data (Tables S7 and S9 in the Supplementary Material). Moreover, uncertain decisions can be easily spotted by small *δ_α,γ_* and 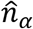 values.

To test the generality of these results, the same procedure was repeated several times by changing the samples of the normal distributions being compared. Hence, it is possible to provide a probability of how often the resulting *θ_α,γ_* would be the same as the one stated in Table 1. Additionally, the presented method has its limitations in the computational time needed to perform MCCV iterations: the more iterations we compute the longer the process will take. Moreover, the accuracy of any estimated *p*(*n*) depends on the sample size *n* = *n_i_* and *p-values*, *p_i_*, that the program can evaluate. Therefore, we also tested the results of the method when the number of iterations *i* and *j* in MCCV is reduced. Overall, a rate between obtaining exactly the same result or a different one under any change of the previous conditions was calculated (Table S8 in the Supplementary Material). The rate is given as a percentage value. The closer the percentage gets to 100, the more robust and general the result will be. We can confirm that the results are most of the time the same as the ones given in Table 1 when *γ* = 5*e*^−06^. The only critical case is the comparison *N*(0, 1) - *N*(0.5, 1) when few *n_i_* points are used to estimate *p*(*n*).

The last procedure was repeated using the real data from the first experiment (study of the effect of Taxol in the cell body and protrusions morphology) (Tables S9 and S10 in the Supplementary Material). Even with more complex and noisier data, the results obtained show that the method is stable and robust. All technical details about these computations are given in the Supplementary Material.

### IMPLEMENTATION DETAILS

We provide a ready-to-use Python code that implements the method explained above, outputs the decision index *θ_α,γ_*, 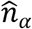, *a* and *c* parameters, and generates the plots shown in the paper : https://github.com/BIIG-UC3M/pMoSS. Together with the implementation, different Jupyter Notebooks with demos and user guidelines are given, making this work accessible from both local machines and Google Colaboratory. The following pseudocode describes the process behind the algorithm. The description of MCCV (`monte_carlo_cross_validation`) is given in the Supplementary Material.

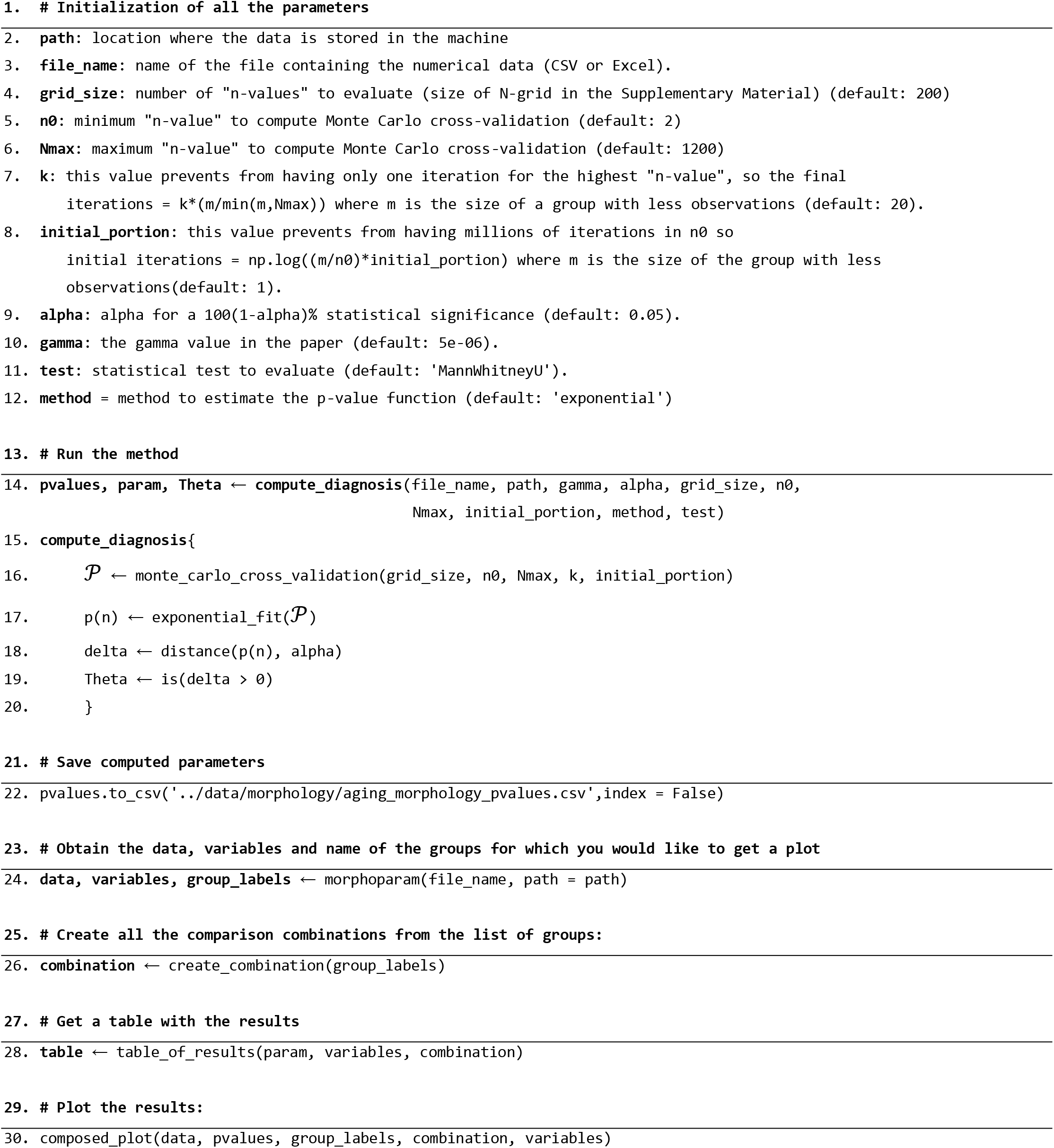

