## Supplementary Material for "Use of the *p-value* as a size-dependent function to address practical differences when analyzing large datasets"

### Supplementary Information

**E. Gómez-de-Mariscal, V. Guerrero, A. Sneider, H. Jayatilaka, J. M. Phillip, D. Wirtz and A. Muñoz-Barrutia**

**Corresponding author: Arrate Muñoz-Barrutia**
****
**Code availability at https://github.com/BIIG-UC3M/pMoSS**

### 1. Technical details

The main motivation of the study is that the *p-value* is no longer useful when working with large datasets as its value tends to zero. In the next section, we demonstrate for the particular cases of the Mann-Whitney U test, (1), and Student's t-test, (2), that indeed, the *p-value* will always tend to zero even when the null hypothesis is almost true and should not be rejected.

**A. *p-values* tend to zero for large sample sizes.** The statistic U of the Mann-Whitney U test, (1), is defined as $\min\{U_1, U_2\}$, where $U_i$ follows the Eq. (1), being $n_i$ the size of the dataset $i$ and $R_i$ its rank sum.

$$U_i = n_1 n_2 + \frac{n_i(n_i+1)}{2} - R_i, \quad i \in \{1,2\}. \tag{1}$$

When $n_i$ are large enough, U follows a normal distribution, (1), with mean and standard deviation values, $\mu_\mathcal{U}$ and $\sigma_\mathcal{U}$ respectively, described by Eq. (2).

$$\mu_\mathcal{U} = \frac{n_1 n_2}{2}, \quad \sigma^2_\mathcal{U} = \frac{n_1 n_2 (n_1 + n_2 + 1)}{12}. \tag{2}$$

Therefore, the main procedure to estimate the *p-value* consists of analyzing the standardized value of U, $z$, defined by

$$z = \frac{\mathrm{U} - \mu_\mathrm{U}}{\sigma_\mathrm{U}}. \tag{3}$$

Replacing the values of $U_i$, $\mu_\mathcal{U}$ and $\sigma^2_\mathcal{U}$ in Eq. (3), we obtain

$$z = \sqrt{12}\left(\frac{n_1 n_2 + \frac{n_1(n_1+1)}{2} - R_1 - \frac{n_1 n_2}{2}}{\sqrt{n_1 n_2 (n_1 + n_2 + 1)}}\right). \tag{4}$$

Since U is defined as the minimum of $U_1$ and $U_2$, we assume for simplicity that $\mathrm{U} = U_1$, but same results hold for $\mathrm{U} = U_2$.

In the worst case scenario, when both datasets are identical and therefore the null hypothesis is true, $R_1 = R_2 = R$. Also, as $n_i$ are assumed to be large enough, we can study the case $n_1 = n_2 = n$. Moreover, due to the hypothesized large sample size, $R_i$ could be upper limited as

$$R \leq \sum_{i=1}^{n} i = \frac{n(n+1)}{2} \qquad \Longrightarrow \qquad z \geq \sqrt{12}\left(\frac{\frac{n^2}{2}}{\sqrt{n^2(2n+1)}}\right) = \frac{n\sqrt{3}}{\sqrt{(2n+1)}}. \tag{5}$$

Finally, the value of $z$ in the limit, when $n$ tends to infinity, is also infinity

$$\lim_{n\to\infty} z \geq \lim_{n\to\infty} \frac{n\sqrt{3}}{\sqrt{(2n+1)}} \longrightarrow \infty \qquad \Longrightarrow \qquad \lim_{n\to\infty} z \longrightarrow \infty. \tag{6}$$

Therefore, *p-value* tends to zero. That is to say, even when we assume that both datasets are equal, the result would be to regret the null hypothesis. Likewise, Student's t-test (2) fails by means of large samples. The statistic $\mathbf{t}$ is defined as follows

$$\mathbf{t} = \frac{\mu_1 - \mu_2}{\sqrt{\frac{S^2}{n_1} + \frac{S^2}{n_2}}}, \tag{7}$$

where $\mu_i$ and $n_i$ correspond to the mean and sample size of the dataset $i \in \{1, 2\}$. Once again, assuming that both $n_i$ are large enough, $n_i = n$ is accepted; $\mathbf{t}$ is directly compared with the Student's t distribution and in the limit of $n$, $t$ tends to infinity (as long as both mean values are not exactly the same). Thus, *p-value* tends to zero and the null hypothesis is rejected.

**B. *p-values* as a function of the sample size.** As it is shown in Fig. 1 of the main manuscript, Fig. 5 in the Materials and Methods, and Section A, the *p-value* depends on the size of the data being evaluated. While this is not a breakthrough, it is one of the pillars in this study. The fact that the *p-value* varies with $n$, allows us to assume that it can be estimated through either a parametric or a non-parametric model. In this case, we fit a parametric model (i.e. the exponential function) to the *p-values* obtained empirically.

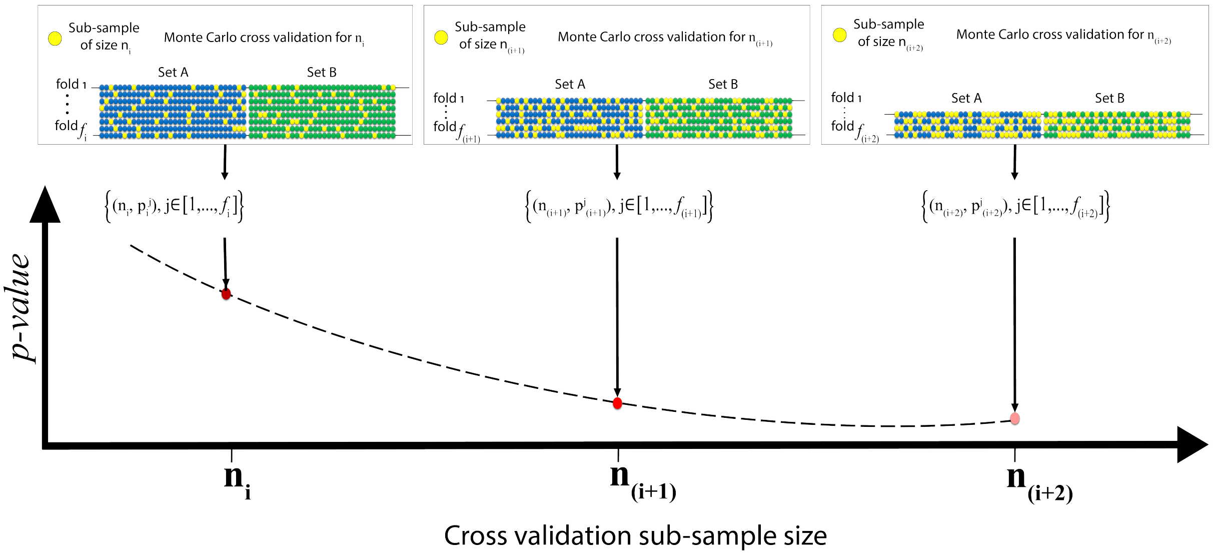

**Fig. S1.** Illustration of the work flow used for the estimation of *p-values* as a function ($p(n)$) of the sample size ($n$). For each possible value of $n$ ($n_i$), Monte Carlo cross validation (MCCV) is performed $f_i$ times. For each fold in the cross validation, two random sub-samples of size $n_i$ are chosen from samples A and B (yellow spheres). Then, a statistical test is applied to obtain a *p-value* ($p_i^j$). The procedure is repeated $f_i$ times; $f_i$ depends on $n_i$ as both samples A and B have to be covered. Thereby, $f_i$ decreases ($f_i > f_{i+1} > f_{i+2}$) as long as $n_i$ gets larger. This procedure is repeated for $n_0$, ..., $n_{i+1}$,$n_{i+2}$, ... until the desired data size ($n_{i\to\infty}$) is reached.

**C. Estimation of *p-values* with Monte Carlo cross validation method.** We propose to model the *p-value* empirically as a data's size dependent function ($p(n)$) by Monte Carlo cross validation (MCCV) with replacement (3). This way, the effect of the sample bias on the *p-value* can be mitigated. The procedure followed to estimate of the *p-values* is illustrated in Figure S1 and the corresponding pseudocode in Algorithm 1. Notice that in Algorithm 1 we estimate $p(n)$ in two different ways, using either a locally weighted scatter plot smoothing (LOWESS) approximation (4) ($p_L$) or and exponential fit ($p_e$). The main reason to do this is that we use a standard curve fitting (LOWESS) to show that $p(n)$ can be expressed as an exponential function.

Aiming to compare two sets of values, $S_A$ and $S_B$, and to determine if there exists statistically significant differences between them, the estimation of the *p-values* is done in pairs (i.e., two sub-samples are compared each time). A range of values needs to be defined for both the sample size and the number of folds in MCCV. These are given by the grids $\mathcal{N}$ and $\mathcal{F}$, respectively.

**Algorithm 1** *p-value* estimation

$N_{max} \leftarrow \min\{|S_A|, |(S_B|\}$

$\mathcal{N} \leftarrow \exp(grid[\log(n_0), \log(n_\infty), gridsize])$
$\mathcal{N} \leftarrow int(\mathcal{N})$

$a \leftarrow N_{max}/k_1 n_0$
$b \leftarrow k_2 N_{max}/n_\infty$
$\mathcal{F} \leftarrow \exp(grid[\log(a), \log(b), gridsize])$
$\mathcal{F} \leftarrow int(\mathcal{F})$

**for** $i$ in $0 : \text{length}(\mathcal{N})$ **do**
  $n_i \leftarrow \mathcal{N}[i]$
  # Start Monte Carlo cross validation:
  **for** $f = 0 : \mathcal{F}[i]$ **do**
    $s_A \leftarrow \text{sample}(S_A, n_i)$
    $s_B \leftarrow \text{sample}(S_B, n_i)$
    $\mathcal{P}_i \leftarrow$ save the *p-value* of $test(s_A, s_B)$
  $\mathcal{P} \leftarrow$ save $mean(\mathcal{P}_i)$
  $\bar{\mathcal{P}} \leftarrow$ save $mean(\mathcal{P}_i)$
$p_L \leftarrow LOWESS(\bar{\mathcal{P}})$
$p_e \leftarrow exponential.fit(\mathcal{P})$

The range for all possible sub-sample sizes ($n$) goes from 2 ($n_0$) to the smallest size between samples $S_A$ and $S_B$ ($N_{max}$). A grid covering all these values for large $N_{max}$, is computationally expensive and redundant. As the *p-value* tends to zero when $n \to \infty$, the most important information is condensed in the smallest values of $n$. So the grid $\mathcal{N}$ follows an exponential distribution from $n_0 = 2$. Similarly, a large enough upper-limit ($n_\infty$) is chosen such that it ensures a fast computation ($n_\infty \ll N_{max}$) and the convergence to zero of $p(n)$. Hence, $\mathcal{N}$ is determined as

$$\mathcal{N} = \{n_i : n_i \in \exp\left(\mathcal{U}\left(\log(n_0), \log(n_\infty)\right)\right)\}, \quad [8]$$

where $\mathcal{U}$ is the uniform distribution that goes from $\log(n_0)$ to $log(n_\infty)$. In MCCV, the number of folds can be extremely large when working with large datasets and a small partition. On the contrary, for a large partition size, the number of folds might decrease dramatically. To compensate for both situations, $\mathcal{F}$ is defined as given below

$$\mathcal{F} = \{f_i : f_i \in \exp\left(\mathcal{U}\left(a, b\right)\right)\}, \text{ where } a = \log\left(\frac{N_{max}}{k_1 n_0}\right), b = \log\left(\frac{k_2 N_{max}}{n_\infty}\right), \quad [9]$$

$\mathcal{U}$ is the uniform distribution, $k_1$ controls the upper-limit on the number of folds for small sub-sample sizes, and $k_2$ controls the lower-limit for large sub-sample sizes. Note that the number of elements in $\mathcal{N}$ and $\mathcal{F}$ are the same.

Finally, for each $n_i$ in $\mathcal{N}$, MCCV is applied to obtain the set of *p-values* defined as

$$\mathcal{P}_i = \left\{p_i^j, j \in [1, ..., f_i]\right\}, \quad [10]$$

being $f_i$ the number of folds in $\mathcal{F}$ that corresponds with the sub-sample size $n_i$ in $\mathcal{N}$.

**D. Assessment of minimum data size needed for statistical significance ($n_\alpha$).** The estimation of the *p-value* function $p(n)$ supports the computation of the minimum data size needed to obtain statistically significant differences ($n_\alpha$). This value is the solution to the equation

$$\alpha = ae^{-cn_\alpha}. \quad [11]$$

As explained in the online methods, the parameters $a$ and $c$ are the result of fitting an exponential function to the empirical values $\mathcal{P}_i$ in Equation 10. Therefore, there exists an intrinsic bias in the estimated values $a$ and $c$. Additionally, the estimation of $p(n)$, and specially, its decay (parameter $c$) can be less precise when the data size is small. (See Figure 2*d* in the main manuscript), so $n_\alpha$ in Equation 11 can be biased.

For this reason, the calculation of a more conservative estimator, $\hat{n}_\alpha$, is strongly recommended

$$\hat{n}_\alpha = arg \min_{n_i} \left\{\left(\overline{p_i} + \sigma_{\overline{p_i}}\right) < \alpha\right\}, \quad [12]$$

where $\overline{p_i}$ represents the mean of $\mathcal{P}_i$ and $\sigma_{\overline{p_i}}$, the mean standard error (SEM), which is included to correct for the variability of the estimated *p-values*. Hence, the estimator of the theoretical value will always be slightly larger

$$\hat{n}_\alpha \geq n_\alpha.$$

However, $\hat{n}_\alpha$ can only be provided when the sample is large enough to cover those $n$ values smaller or equal to $\hat{n}_\alpha$. For this reason, whenever the data is not large enough, the theoretical value $n_\alpha$ in Equation 11 is also given. In these cases, even if $n_\alpha$ and $\Theta_{\alpha,\gamma}$ might be slightly deviated, they still serve as an indicator of the existence of statistical significance.

### 2. Experimental data

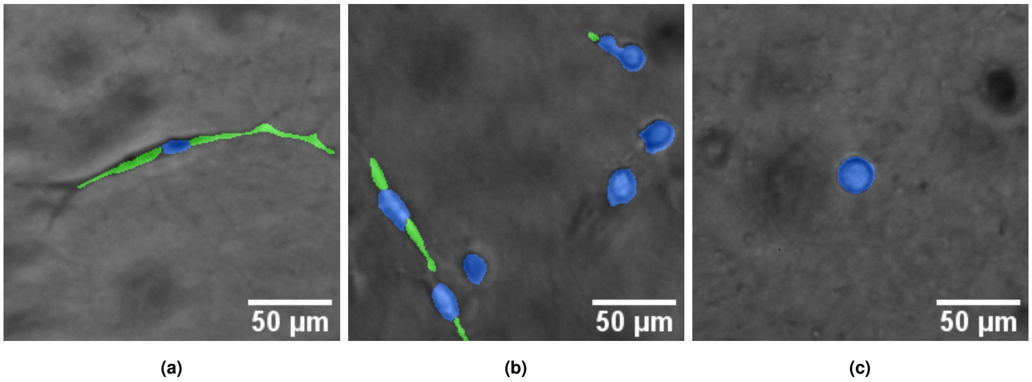

**Fig. S2.** Segmentation of phase contrast microscopy images of cancer cells (MDA-MB-231) embedded in a 3D collagen Type I matrix. Cell bodies are labeled in blue and cell protrusions in green. Images of (a) control cells and cells treated at (b) 1 nM and (c) 50 nM Taxol were acquired with a 10 x magnification objective.

**A. Experiment 1: Drug analysis on phase contrast microscopy data.** Image processing analysis provided the necessary information to distinguish the cellular body and protrusions of each cell in the videos (Figure S2). Hence, we got eight different measurements: cell body size ($C_S$), cell body perimeter ($C_P$), cell body roundness ($C_R$), cell with at least one protrusion ($P_b$), protrusion size ($P_S$), protrusion perimeter ($P_P$), protrusion length ($P_L$) and protrusion diameter ($P_D$). Each of the morphological measurements is given in microns. Table S1 contains the complete list of variables.

**Table S1. List of computed variables. C: continuous variable. B: binary variable.**

| Cell body | | | Cell protrusions | | |
|---|---|---|---|---|---|
| Feature | Name | Type | Feature | Name | Type |
| Area ($\mu m^2$) | $C_S$ | Categorical | Area ($\mu m^2$) | $P_S$ | Categorical |
| Perimeter ($\mu m$) | $C_P$ | Categorical | Perimeter ($\mu m$) | $P_P$ | Categorical |
| Roundness | $C_R$ | Categorical | Length ($\mu m$) | $P_L$ | Categorical |
| Protrusions | $P_b$ | Binary | Diameter ($\mu m$) | $P_D$ | Categorical |

In Figure S3, the distribution of the the variables used in the analysis of cellular shape is shown. The cellular body changes with the amount of Taxol used to treat cells. When they are treated at 50 nM Taxol, the cellular body is biger and more rounded (Figure S3a). Besides, this same treatment prevents cells from producing long and thick protrusions (Figure S3b).

None of the continuous variables presented in Table S1 follows a normal distribution, so the comparison was carried out by the Mann-Whitney U-test (1). $P_b$ was a binary variable (Figure S3c) to distinguish protruding cells (value one). Therefore, it was analyzed by means of Pearson $\chi^2$-test for categorical data (5).

As per the number of observations reported in Table S2 and following methodology guidelines, we set $\mathcal{P}_i$ with $n_0 = 2$ and $n_\infty = 2500$. $\mathcal{N}$ and $\mathcal{F}$ were set to have 190 points. The number of folds $\mathcal{F}$ described in the Supplementary Material, was computed using $k_1 = 1$, $k_2 = 20$ and $N_{max} = 11037$. These values were chosen to have a reasonable number of permutations for both small and large sample sizes (6.000 permutations when $n_0 = 2$, and 90 when $n_{190} = 2500$, respectively). Table S3 contains the estimated coefficients $a$ and $c$ of the exponential curve ($ae^{-cn}$) for each of the variables we analyzed and each pair of comparisons (Control - 1 nM Taxol, Control - 50 nM Taxol, and 1 nM - 50 nM Taxol).

| Treatment group | Cell body | Cell protrusions |
|---|---|---|
| Control | 77,700 | 45,871 |
| 1 nM Taxol | 74,713 | 42,798 |
| 50 nM Taxol | 46,162 | 11,037 |

**Table S2. Number of observations (cell body and their protrusions -if present-) per treatment group.**

Figures S4, S5 and S6 show the shape of each of the exponential curves that result with the coefficients in Table S3. To determine whether Taxol has a significant effect in cell's morphology, $\Theta_{\alpha,\gamma}$ was chosen such that $\alpha = 0.05$ (95% of statistical significance) and $\gamma = 5 \cdot 10^{-6}$.

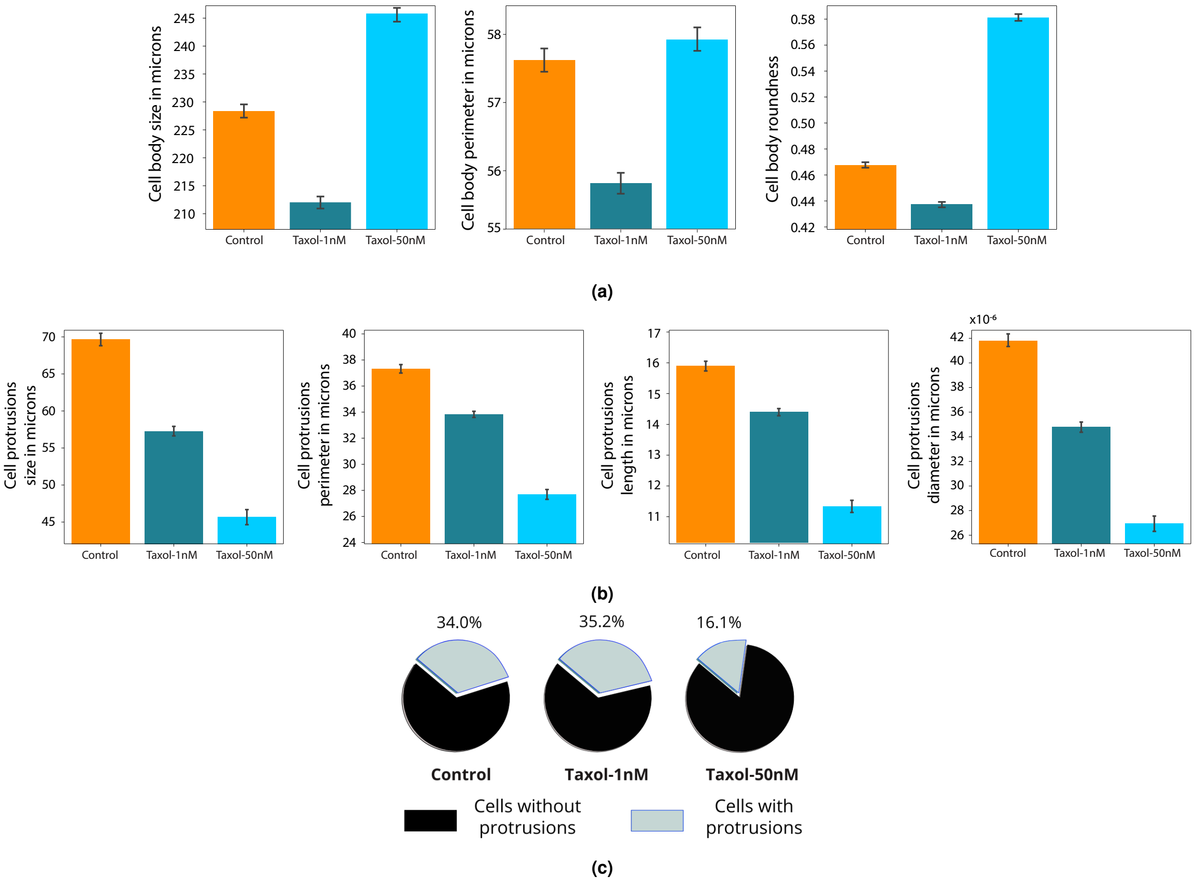

**Fig. S3.** Quantitative variables used to measure (a) cell bodies and (b) cellular protrusions morphology, and (c) the ratio of cells with and without protrusions for the three different treatment groups (control, $1$ nM Taxol, $50$ nM Taxol). Error bars in (a) and (b) correspond to the confidence interval at $99$ %

When comparing the control group and 1 nM Taxol, there are not statistically significant differences in cell body morphology: the curve $p(n)$ of any cell body feature decreases slowly, i.e. $\hat{n}_\alpha$ and $n_\gamma$ are large and $\Theta_{\alpha,\gamma} = 0$ (Table S3 and Figure S4a). On the other hand, cells at 50 nM Taxol have a significantly higher roundness index and bigger cellular body: when comparing control vs. 50 nM Taxol or 1 nM vs. 50 nM Taxol the curves corresponding to $C_R$ and $C_S$ decrease rapidly, i.e. $\hat{n}_\alpha$ and $n_\gamma$ are small, and $\Theta_{\alpha,\gamma} = 1$ (Table S3 and Figures S4b and S4c, respectively). For $C_P$, it is also possible to appreciate some differences when comparing 1 nM with 50 nM Taxol group, i.e. $\Theta_{\alpha,\gamma} = 1$ (Table S3). Namely, the blue curve shown in Figure S4c decreases faster than those in Figures S4a and S4b.

Similar results are obtained when the morphology of cellular protrusions is evaluated (Table S3 and Figure S5). While Taxol at 1 nM does not change their morphology ($\Theta_{\alpha,\gamma} = 0$ in Table S3 and Figure S5a), the effect of Taxol at 50 nM is much larger ($\Theta_{\alpha,\gamma} = 1$ in Table S3, Figures S4b and S4c) .

Usually, when a categorical variable such as $P_b$ is analyzed, the input of a statistical test is a percentage rather than the raw data. Hence, when there is no statistical significance, the $p(n)$ function shoots up, as for instance in Figure S6a. However, when there exist statistical differences, $p(n)$ decreases and it is possible to analyze its decay, as in Figures S6b-c. With all, we can say that the formation of protrusions is inhibited when 50 nM Taxol are administered: there is a significant reduction in the number of cells that form protrusions and their protrusions are smaller (shorter and thinner) (S3, Figures S3b and S3c).

**B. Experiment 2: Drug analysis on flow cytometry data.** Flow cytometry is a technique that generates a large amount of data for each experiment. Consequently, any statistical test for groups comparison results in a vanishing *p-value*. To avoid that situation, practitioners tend to reduce the data to a single, representative measure for subject. For instance, Khoury *et al.* (6) acquired fluorescence intensity data from 6 different subjects and compute the median fluorescence intensity (MFI) for each of them. So, the statistical test is just applied on the 6 MFI values. However, our proposal of estimating the *p-value* as a function of the sample size enables to incorporate in the test the information given by the whole dataset and take into consideration the deviation and bias present in the data.

To illustrate the proposed procedure, we analyzed the flow cytometry data provided in (7) to determine the transcriptional changes induced by the *in vivo* exposure of human eosinophils to glucocorticoids. Khoury *et al.* (6) studied eosinophil surface proteins after being exposed to glucocorticoids and demonstrated that this exposure causes the apoptosis of human eosinophils

| Variables | Cell body size ($C_S$) | | | | Cell body perimeter ($C_P$) | | | | Cell body roundness ($C_R$) | | | | Cell with protrusions ($P_b$) | | | |
|---|---|---|---|---|---|---|---|---|---|---|---|---|---|---|---|---|
| Comparison | $a$ | $c$ | $\hat{n}_\alpha$ | $\Theta_{\alpha,\gamma}$ | $a$ | $c$ | $\hat{n}_\alpha$ | $\Theta_{\alpha,\gamma}$ | $a$ | $c$ | $\hat{n}_\alpha$ | $\Theta_{\alpha,\gamma}$ | $a$ | $c$ | $\hat{n}_\alpha$ | $\Theta_{\alpha,\gamma}$ |
| C - 1T | 0.258 | 0.0026 | 670 | **0** | 0.258 | 0.0017 | 1160 | **0** | 0.259 | 0.0029 | 617 | **0** | 0.435 | -0.0005 | $\infty$ | **0** |
| C - 50T | 0.263 | 0.0075 | 250 | 1 | 0.256 | 0.0014 | 1331 | **0** | 0.282 | 0.0400 | 47 | 1 | 0.198 | 0.0345 | 42 | 1 |
| 1T - 50T | 0.272 | 0.0216 | 83 | 1 | 0.264 | 0.0072 | 257 | 1 | 0.292 | 0.0648 | 29 | 1 | 0.195 | 0.0351 | 41 | 1 |
| Variables | Protrusions size ($P_S$) | | | | Protrusions perimeter ($P_P$) | | | | Protrusions length ($P_L$) | | | | Protrusions diameter ($P_D$) | | | |
| Comparison | $a$ | $c$ | $\hat{n}_\alpha$ | $\Theta_{\alpha,\gamma}$ | $a$ | $c$ | $\hat{n}_\alpha$ | $\Theta_{\alpha,\gamma}$ | $a$ | $c$ | $\hat{n}_\alpha$ | $\Theta_{\alpha,\gamma}$ | $a$ | $c$ | $\hat{n}_\alpha$ | $\Theta_{\alpha,\gamma}$ |
| C - 1T | 0.250 | 0.0031 | 563 | 1 | 0.248 | 0.0019 | 754 | **0** | 0.251 | 0.0011 | 1695 | **0** | 0.251 | 0.0023 | 707 | **0** |
| C - 50T | 0.246 | 0.0221 | 75 | 1 | 0.241 | 0.0276 | 58 | 1 | 0.250 | 0.0289 | 58 | 1 | 0.250 | 0.0248 | 68 | 1 |
| 1T - 50T | 0.250 | 0.0100 | 170 | 1 | 0.256 | 0.0175 | 98 | 1 | 0.255 | 0.0211 | 80 | 1 | 0.247 | 0.0134 | 127 | 1 |

**Table S3. Parameters of the exponential function $ae^{-cn}$ and estimated minimum size $\hat{n}_\alpha$ for each of the analyzed variables. C: control, $1$T: $1$ nM Taxol and $50$T: $50$ nM Taxol.** $a \in [0.195, 0.435]$, $c \in \left[-5 \cdot 10^{-4}, 0.0648\right]$, $\alpha = 0.05$ **and** $\gamma = 5 \cdot 10^{-6}$.

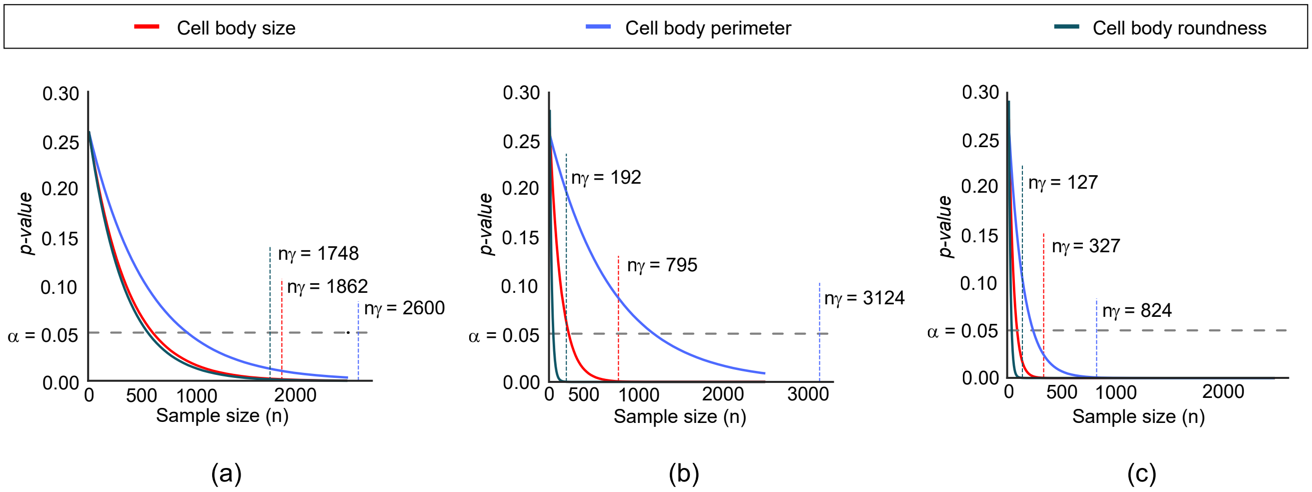

**Fig. S4.** Results obtained for cell body morphology when control, $1$ nM Taxol and $50$ nM Taxol groups are compared as (a) control vs. $1$ nM Taxol, (b) control vs. $50$ nM Taxol and (c) $1$ nM vs. $50$ nM Taxol. Vertical lines correspond to the convergence point $n_\gamma$ with $\gamma = 5 \cdot 10^{-6}$.

(eosinopenia) once they migrate out of the blood circulation.

While they performed an extensive analysis, we have focused our study on the data related to the chemokine receptor gene CXCR4. In particular, the expression of CXCR4 on the surface of human eosinophils after being exposed for 2 hours to vehicle, 20 mcg/dL and 200 mcg/dL of Methylprednisolone (MP). After filtering the raw data to discard noise and debris, we got clean distributions to analyze (Figure 2b, right). The *p-value* curves computed for pair group comparisons were the result of applying Mann-Whitney U statistical tests following the proposed procedure. Then, the exponential curves ($ae^{-cn}$) were fitted (Figure 2b, left). The parameters configuration was $n_0 = 2$, $\mathcal{N}$ and $\mathcal{F}$ had 200 points, $k_1 = 1$, $k_2 = 20$, $\alpha = 0.05$, and $\gamma = 5 \cdot 10^{-6}$. As the number of data points was lower than $1,000$, $n_\infty$ and $N_{max}$ where chosen to be the minimum number of points for each group pair being compared. The results are summarized in Table S4. Our results are similar to those in (6), in the sense that we also find a differential expression of CXCR4 when eosinophils are exposed to glucocorticoids.

**C. Experiment 3: Cellular age characterization by means of biomolecular and biophysical properties.** The data extracted from Phillip *et al.* in (8) was analysed with the following parameter configuration: for each of the stated variables, the data belonging to the group of 2 years-old was compared with the data from $\{3, 9, 16, 29, 35, 55, 65, 85, 92\}$ and $\{3, 9, 16, 29, 35, 55, 65, 85, 96\}$ years-old human donors to test cell motility and morphology, respectively. For each pair of groups, the distribution of the $n$-dependent *p-values* was obtained using the Mann-Whitney U statistical test. Then, the parameters of the exponential function ($ae^{-cn}$) were fitted. The parameter configuration was $n_0 = 2$, $\mathcal{N}$ and $\mathcal{F}$ had 200 points, $k_1 = 1$, $k_2 = 20$, $\alpha = 0.05$, and $\gamma = 5 \cdot 10^{-6}$. As the number of data points was lower than 1000, $n_\infty$ and $N_{max}$ where chosen to be the minimum number of points of each pair of groups being compared. The results for cell motility and cell morphology are summarized in Tables S5 and S6, respectively.

### 3. Test of robustness

The variability in the statistical significance of the results caused by the selection of the parameter $\gamma$ and the grid sizes $\mathcal{N}$ and $\mathcal{F}$ are characterized in this section. The method is first tested using theoretical distributions and then, using the real data from Experiment 1.

| Measures | CXCR4 surface expression | | | | |
|---|---|---|---|---|---|
| Comparison | $a$ | $c$ | $\hat{n}_\alpha$ | $n_\alpha$ | $\Theta_{\alpha,\gamma}$ |
| Vehicle - MP $20$ mcg/dL | 0.286 | 0.048 | 40 | 36 | 1 |
| Vehicle - MP $200$ mcg/dL | 0.286 | 0.049 | 37 | 35 | 1 |
| MP $20$ mcg/dL - MP $200$ mcg/dL | 0.256 | 1.69·10$^{-4}$ | - | 9680 | 0 |

**Table S4. Parameters of the exponential function $ae^{-cn}$ for the differential expression of CXCR4, theoretical minimum size ($n_\alpha$) and its estimator ($\hat{n}_\alpha$) for a $95\%$ ($\alpha = 0.05$) of statistical significance, and decision index $\Theta_{\alpha,\gamma}$, for $\gamma = 5 \cdot 10^{-6}$. MP: Methylprednisolone**

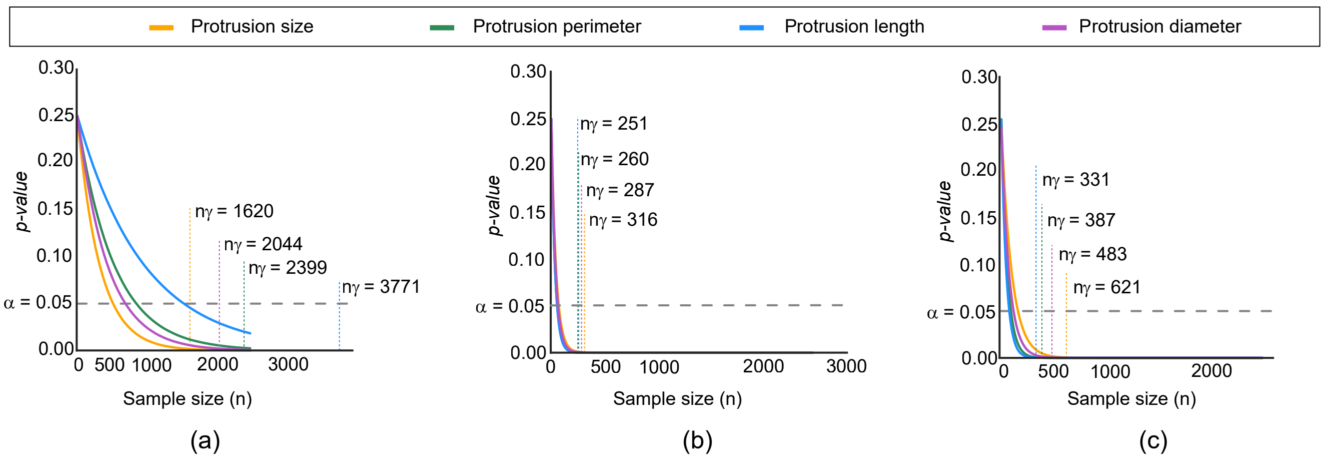

**Fig. S5.** Graphical illustration of the results obtained for cell protrusions morphology when (a) control and 1 nM Taxol, (b) control and 50 nM Taxol and (c) 1 nM Taxol and 50 nM Taxol groups are compared. Vertical lines correspond to the convergence point $n_\gamma$ with $\gamma = 5 \cdot 10^{-6}$.

**A. Test of robustness on theoretical data.** We simulated normal distributions to test the method in a theoretical scenario: $\mathcal{N}(0,1)$ was compared with $\mathcal{N}(0.01,1)$, $\mathcal{N}(0.1,1)$, $\mathcal{N}(0.25,1)$, $\mathcal{N}(0.5,1)$, $\mathcal{N}(0.75,1)$, $\mathcal{N}(1,1)$, $\mathcal{N}(2,1)$ and $\mathcal{N}(3,1)$. For the most similar cases such as $\mathcal{N}(0,1)$ vs. $\mathcal{N}(0.01,1)$, or $\mathcal{N}(0,1)$ vs. $\mathcal{N}(0.1,1)$, it is expected to obtain $\Theta_{\alpha,\gamma} = 0$. While for the most different distributions such as $\mathcal{N}(0,1)$ vs. $\mathcal{N}(2,1)$, or $\mathcal{N}(0,1)$ vs. $\mathcal{N}(3,1)$, $\Theta_{\alpha,\gamma} = 1$.

Theoretically, an optimal grid $\mathcal{N}$ would be the one that covers the values from $n_0 = 2$ to $n_\infty = N_{max}$. This set up entails an extremely large amount of computations, while it suffices a value $n_\infty \approx 1000$ to understand what is the tendency of the data. If $p(n)$ converges to zero when $n > 1,000$, then it can be assumed that $p(n)$ does not represent a statistical significant case. Hence, $n_\infty = 2,500$ is large enough for the implementation of the method. As the *p-values* for very small samples are especially unstable and small samples are not representative of any real scenario, the minimum value $n_0$ can be increased. The number of permutations for each $n_i$ can be decreased as well: while the amount of data to analyze may be infinite, it is enough to study a certain limited number of different data subsamples to approach a realistic scenario. Hence, to test the robustness of the proposed method, we set grids $\mathcal{N}$ and $\mathcal{F}$ using $n_0 = 20$, $n_\infty = 2,500$, $N_{max} = 10,000$, $k_1 = 1$ and $k_2 = 20$ in Equations 8 and 9, respectively. Both $\mathcal{N}$ and $\mathcal{F}$ were configured to have a size of 200 points. Thus, MCCV is repeated 200 times. See Figure S1 for the workflow.

With this grid parameters, for $\gamma$ in the set $\{2.5 \cdot 10^{-6}, 5 \cdot 10^{-6}, 5 \cdot 10^{-5}, 5 \cdot 10^{-4}\}$, we run the pipeline to evaluate the effect of $\gamma$ value on the rejection of the null hypothesis of the Mann-Whitney U statistical test (Table S7). The results obtained help to assess the most suitable $\gamma$ value. Specifically, the decision about $\gamma$ relies on the result obtained for the comparison between $\mathcal{N}(0,1)$ and $\mathcal{N}(0.25,1)$: while the distance $\delta_{\alpha,\gamma}$ for $\gamma = 5 \cdot 10^{-6}$ and $\gamma = 5 \cdot 10^{-5}$ expresses the same ($\delta_{\alpha,\gamma} = \pm 5.84$), the minimum data size needed to observe statistically significance differences is low enough as to reject the null hypothesis, i.e. $\hat{n}_\alpha = 186$ and $\Theta_{\alpha,\gamma} = 1$. Hence, the value chosen for the following simulations and for the real data is $\gamma = 5 \cdot 10^{-6}$. (Table S7).

To test the computational limitations of the method, we evaluated the value $\Theta_{0.05, 5 \cdot 10^{-6}}$ reducing $\mathcal{N}$ and $\mathcal{F}$: $\mathcal{N}$ was chosen to be a grid of size 10, 20, 50, 100, 150 or 200 points and the values in $\mathcal{F}$ were reduced by a factor of 1/2, 1/3, 1/5 and 1/10 (i.e., each of the values in the original $\mathcal{F}$ was multiplied by this fraction). The experiment was repeated 100 times on each of the setups, so the probability of obtaining exactly the same $\Theta_{0.05, 5 \cdot 10^{-6}}$ (Table S7) and the stability of the method could be evaluated. The information given in Table S8 lets the assessment of (1) the size of $\mathcal{N}$ and (2) the number of folds in $\mathcal{F}$. In most cases, the probability obtained was 100%, which shows that the final results are very stable. When $N(0,1)$ and $N(0.5,1)$ were compared with small grid parameters, this probability decreased slightly to $89-96\%$ (Table S8). In conclusion, the number of computations could be considerably reduced, for the example, to $\mathcal{N} = 50$ and $\mathcal{F} = 0.2\mathcal{F}$).

**B. Test of robustness on real data.**

***B.1. The* p-value *can be estimated by an exponential function..*** Repeating the procedure followed with simulation of normal distributions, we verify that the condition for $p(n)$ being exponential is satisfied again: in Figures S7a-c all LOWESS fittings have exponential shapes, and in Figures S7d-f the quotient $p'(n)/p(n)$ of LOWESS fits are constant.

***B.2. Robustness of the convergence threshold and required computational load.*** The distribution of real data is more complex than the typical Gaussian distributions due to the presence of noise, large deviation of the data or leverage points. Following the same procedure as in Section A, we tested the reliability of the proposed method using the data we extracted from the microscopy images (Experiment 1, main manuscript). We evaluated both, the effect of varying the convergence threshold $\gamma$ and the required computational load. Looking at Tables S3 and S9, it can be appreciated once again that $\gamma = 5 \cdot 10^{-6}$

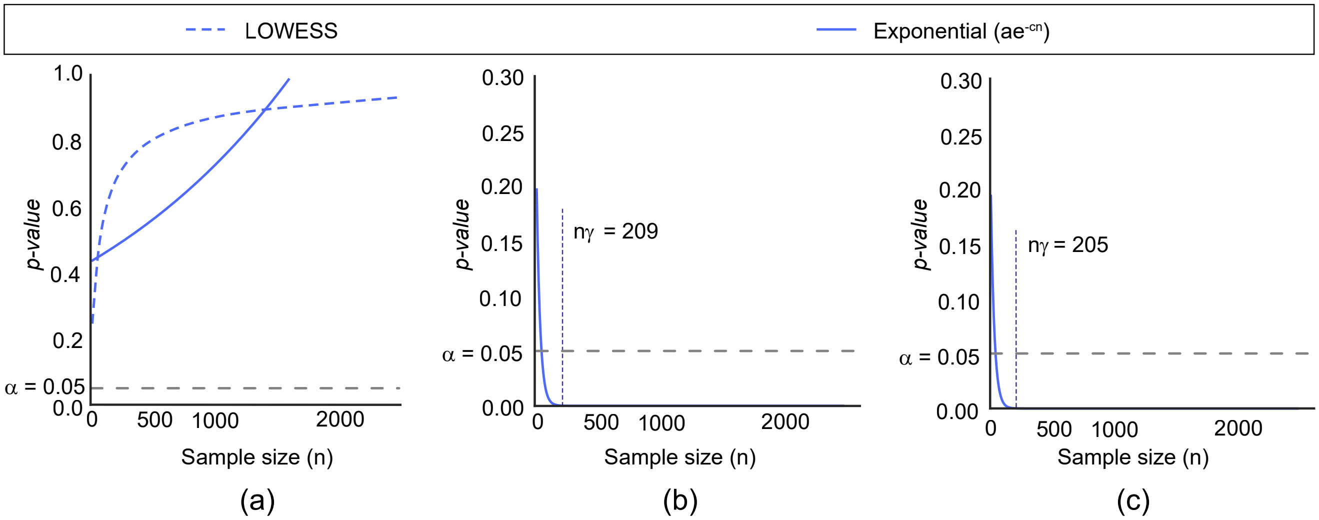

**Fig. S6.** Estimation of $p(n)$ with $\chi^2$ test for contingency tables when comparing cells (not) having at least one protrusion (binary variable cell with protrusions: $P_b$) for (a) control and $1$ nM Taxol groups, (b) control and $50$ nM Taxol groups, and (c) $1$ nM and $50$ nM Taxol groups. In the leftmost plot, both locally weighted scatter plot smoothing (LOWESS) and exponential fit of estimated *p-values* are shown. Vertical lines correspond to the convergence point $n_\gamma$ with $\gamma = 5 \cdot 10^{-6}$.

is a good value for the convergence threshold. Smaller values of $\gamma$ result in the rejection of the null hypothesis for cases in which $\hat{n}_\alpha > 1000$ as cellular protrusions length. Similarly, when $\gamma = 5 \cdot 10^{-5}$, there are cases as cell body roundness for which $\hat{n}_\alpha < 100$ and $\Theta_{\alpha,\gamma} = 0$. Therefore, once again, $\gamma = 5 \cdot 10^{-6}$ seems to be an appropriate value to measure statistical significance at $\alpha = 0.05$ significance level (Table S9).

When the grid $\mathcal{N}$ is large enough and $\mathcal{F}$ has large numbers, the result is completely stable (Table S10). However, when these values are dramatically reduced (for example, $\mathcal{N} = 10$, $\mathcal{F} = 0.02\mathcal{F}_0$, $\mathcal{F} = 0.01\mathcal{F}_0$), the reproducibility of the results may degrade. This fact is specially evident when the variables are noisy as in the case of those that measure cellular protrusions morphology ($P_S$, $P_P$, $P_L$ and $P_D$), being the noisier the protrusions size and perimeter. In summary, while large grid parameters ensure stable results, with the information given in Tables S8 and S10, seems reasonable to reduce the number of computations to $\mathcal{N} \geq 50$ and $\mathcal{F} \geq 0.2\mathcal{F}$.

| Measures | MSD-6min | | | | | MSD-60 min | | | | | Persistence primary axis | | | | |
|---|---|---|---|---|---|---|---|---|---|---|---|---|---|---|---|
| Comparison | $a$ | $c$ | $\hat{n}_\alpha$ | $n_\alpha$ | $\Theta_{\alpha,\gamma}$ | $a$ | $c$ | $\hat{n}_\alpha$ | | $\Theta_{\alpha,\gamma}$ | $a$ | $c$ | $\hat{n}_\alpha$ | $n_\alpha$ | $\Theta_{\alpha,\gamma}$ |
| A02 - A03 | 0.296 | 0.038 | 46 | 46 | 1 | 0.276 | 0.009 | - | 187 | 1 | 0.287 | 0.012 | - | 145 | 1 |
| A02 - A09 | 0.270 | 0.001 | - | 1333 | 0 | 0.295 | 0.000 | - | $4.72 \cdot 10^{17}$ | 0 | 0.276 | 0.002 | - | 896 | 0 |
| A02 - A16 | 0.289 | 0.014 | - | 125 | 1 | 0.262 | 0.001 | - | 1955 | 0 | 0.310 | 0.084 | 23 | 21 | 1 |
| A02 - A29 | 0.394 | 0.217 | 12 | 9 | 1 | 0.357 | 0.161 | 13 | 12 | 1 | 0.287 | 0.013 | - | 137 | 1 |
| A02 - A35 | 0.315 | 0.155 | 12 | 11 | 1 | 0.335 | 0.123 | 18 | 15 | 1 | 0.274 | 0.002 | - | 752 | 0 |
| A02 - A55 | 0.286 | 0.018 | - | 99 | 1 | 0.283 | 0.032 | 52 | 54 | 1 | 0.286 | 0.000 | - | $4.99 \cdot 10^{15}$ | 0 |
| A02 - A65 | 0.306 | 0.032 | 48 | 56 | 1 | 0.345 | 0.113 | 15 | 17 | 1 | 0.330 | 0.097 | 19 | 19 | 1 |
| A02 - A85 | 0.282 | 0.008 | - | 229 | 1 | 0.303 | 0.040 | 46 | 45 | 1 | 0.315 | 0.050 | 34 | 36 | 1 |
| A02 - A92 | 0.345 | 0.189 | 12 | 10 | 1 | 0.386 | 0.228 | 10 | 8 | 1 | 0.331 | 0.079 | 27 | 24 | 1 |
| Measures | Persistence secondary axis | | | | | Diffusivity primary axis | | | | | Diffusivity secondary axis | | | | |
| Comparison | $a$ | $c$ | $\hat{n}_\alpha$ | $n_\alpha$ | $\Theta_{\alpha,\gamma}$ | $a$ | $c$ | $\hat{n}_\alpha$ | $n_\alpha$ | $\Theta_{\alpha,\gamma}$ | $a$ | $c$ | $\hat{n}_\alpha$ | $n_\alpha$ | $\Theta_{\alpha,\gamma}$ |
| A02 - A03 | 0.276 | 0.001 | - | 1399 | 0 | 0.301 | 0.000 | - | 1.47E+17 | 0 | 0.290 | 0.034 | 52 | 51 | 1 |
| A02 - A09 | 0.266 | 0.001 | - | 3023 | 0 | 0.275 | 0.006 | - | 289 | 1 | 0.279 | 0.051 | 32 | 33 | 1 |
| A02 - A16 | 0.295 | 0.069 | 27 | 25 | 1 | 0.293 | 0.030 | 56 | 58 | 1 | 0.268 | 0.004 | - | 443 | 0 |
| A02 - A29 | 0.301 | 0.019 | - | 94 | 1 | 0.335 | 0.078 | 27 | 24 | 1 | 0.354 | 0.160 | 15 | 12 | 1 |
| A02 - A35 | 0.281 | 0.014 | - | 126 | 1 | 0.312 | 0.061 | 30 | 29 | 1 | 0.312 | 0.117 | 19 | 15 | 1 |
| A02 - A55 | 0.284 | 0.015 | - | 116 | 1 | 0.285 | 0.016 | - | 109 | 1 | 0.284 | 0.008 | - | 213 | 1 |
| A02 - A65 | 0.294 | 0.030 | - | 58 | 1 | 0.327 | 0.086 | 24 | 21 | 1 | 0.312 | 0.100 | 15 | 18 | 1 |
| A02 - A85 | 0.287 | 0.027 | 54 | 65 | 1 | 0.304 | 0.051 | 33 | 35 | 1 | 0.269 | 0.004 | - | 397 | 1 |
| A02 - A92 | 0.293 | 0.016 | - | 112 | 1 | 0.380 | 0.217 | 11 | 9 | 1 | 0.295 | 0.025 | - | 71 | 1 |
| Measures | Total diffusivity | | | | | Anisotropy | | | | | | | | | |
| Comparison | $a$ | $c$ | $\hat{n}_\alpha$ | $n_\alpha$ | $\Theta_{\alpha,\gamma}$ | $a$ | $c$ | $\hat{n}_\alpha$ | $n_\alpha$ | $\Theta_{\alpha,\gamma}$ | | | | | |
| A02 - A03 | 0.284 | 0.000 | - | $8.70 \cdot 10^{18}$ | 0 | 0.291 | 0.024 | 69 | 72 | 1 | | | | | |
| A02 - A09 | 0.280 | 0.000 | - | $1.68 \cdot 10^{19}$ | 0 | 0.307 | 0.075 | 26 | 24 | 1 | | | | | |
| A02 - A16 | 0.297 | 0.036 | 40 | 48 | 1 | 0.293 | 0.000 | - | $1.16 \cdot 10^{15}$ | 0 | | | | | |
| A02 - A29 | 0.360 | 0.122 | 17 | 16 | 1 | 0.290 | 0.016 | - | 106 | 1 | | | | | |
| A02 - A35 | 0.330 | 0.093 | 22 | 20 | 1 | 0.281 | 0.014 | - | 120 | 1 | | | | | |
| A02 - A55 | 0.276 | 0.006 | - | 269 | 1 | 0.309 | 0.043 | 40 | 42 | 1 | | | | | |
| A02 - A65 | 0.330 | 0.105 | 19 | 18 | 1 | 0.266 | 0.000 | - | 35475 | 0 | | | | | |
| A02 - A85 | 0.306 | 0.052 | 33 | 34 | 1 | 0.293 | 0.015 | - | 120 | 1 | | | | | |
| A02 - A92 | 0.379 | 0.201 | 11 | 10 | 1 | 0.359 | 0.104 | 20 | 19 | 1 | | | | | |

**Table S5. Parameters of the exponential function $ae^{-cn}$ for cell motility, theoretical minimum size ($n_\alpha$) and estimated one ($\hat{n}_\alpha$) for a $95\%$ ($\alpha = 0.05$) of statistical significance, and decision index $\Theta_{\alpha,\gamma}$ , for $\gamma = 5 \cdot 10^{-6}$.**

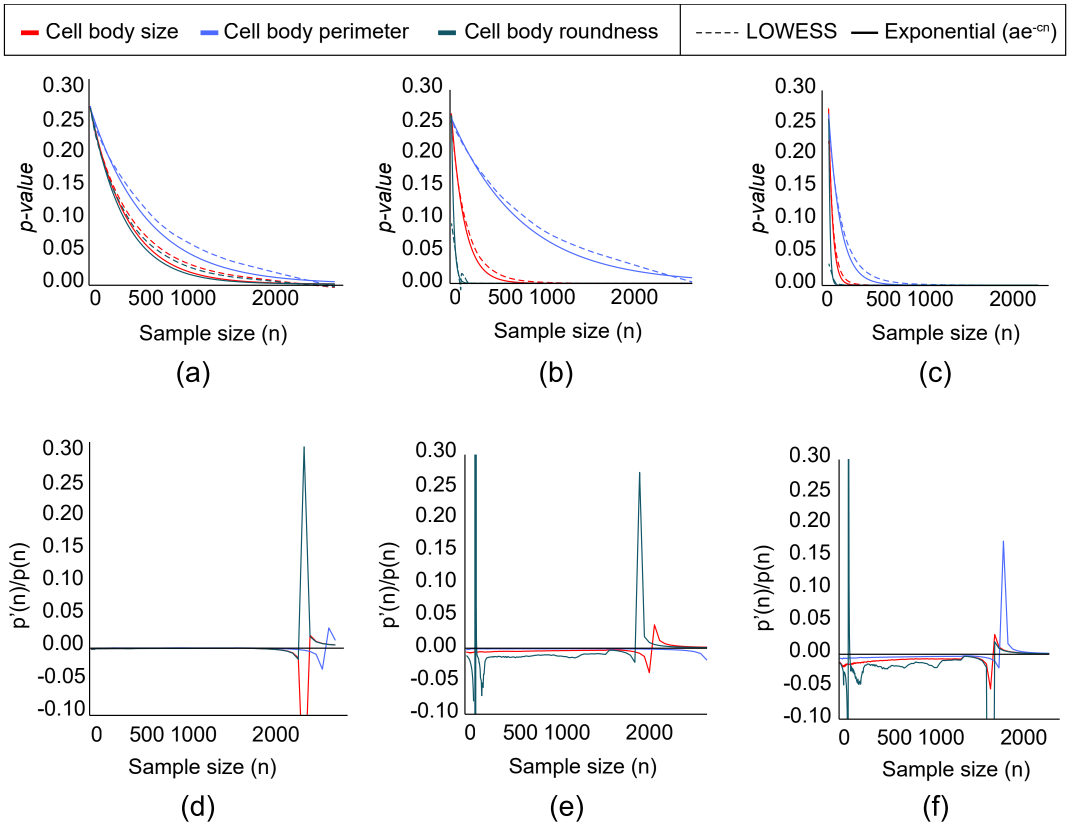

**Fig. S7.** Curve fitting. Three different simulations of $p(n)$ are computed with the data obtained from the experimental test. Cell body size, perimeter and roundness of three different cell groups where compared by means of the Mann-Whitney U statistical test and Monte Carlo cross validation: (a) and (d) control cells versus cells treated with Taxol at $1$ nM; (b) and (e) control cells versus cells treated with Taxol at $50$ nM and (c) and (f) cells treated with Taxol at $1$ nM and $50$ nM. To each of the mean *p-value* sets (i.e., the output of Monte Carlo cross validation), a locally weighted scatter plot smoothing (LOWESS) (4) curve was fit to get the initial shape of $p(n)$. Likewise, an exponential function was fit to all the *p-values* obtained in each fold of the Monte Carlo cross validation (before averaging). Both the LOWESS and exponential curves are shown in (a), (b) and (c). The quotient between each LOWESS curve and its differential is shown in (d), (e) and (f). The constant quotients and the accurate exponential fits show empirically that the $p(n)$ functions have an exponential nature.

| Measures | Size (pixels) | | | | | Perimeter (pixels) | | | | | Long axis length (pixels) | | | | |
|---|---|---|---|---|---|---|---|---|---|---|---|---|---|---|---|
| Comparison | $a$ | $c$ | $\hat{n}_\alpha$ | $n_\alpha$ | $\Theta_{\alpha,\gamma}$ | $a$ | $c$ | $\hat{n}_\alpha$ | $n_\alpha$ | $\Theta_{\alpha,\gamma}$ | $a$ | $c$ | $\hat{n}_\alpha$ | $n_\alpha$ | $\Theta_{\alpha,\gamma}$ |
| A02 - A03 | 0.267 | 0.002 | 509 | 688 | 0 | 0.271 | 0.009 | 203 | 183 | 1 | 0.276 | 0.022 | 76 | 76 | 1 |
| A02 - A09 | 0.264 | 0.002 | - | 923 | 0 | 0.260 | 0.000 | - | 22578 | 0 | 0.274 | 0.008 | - | 201 | 1 |
| A02 - A16 | 0.320 | 0.149 | 13 | 12 | 1 | 0.340 | 0.210 | 10 | 9 | 1 | 0.477 | 0.459 | 6 | 4 | 1 |
| A02 - A29 | 0.312 | 0.129 | 16 | 14 | 1 | 0.313 | 0.121 | 16 | 15 | 1 | 0.325 | 0.144 | 14 | 12 | 1 |
| A02 - A35 | 0.311 | 0.106 | 19 | 17 | 1 | 0.295 | 0.084 | 21 | 21 | 1 | 0.316 | 0.133 | 16 | 13 | 1 |
| A02 - A55 | 0.313 | 0.112 | 18 | 16 | 1 | 0.323 | 0.136 | 13 | 13 | 1 | 0.364 | 0.243 | 10 | 8 | 1 |
| A02 - A65 | 0.326 | 0.137 | 16 | 13 | 1 | 0.373 | 0.270 | 8 | 7 | 1 | 0.401 | 0.335 | 7 | 6 | 1 |
| A02 - A85 | 0.330 | 0.167 | 13 | 11 | 1 | 0.431 | 0.401 | 6 | 5 | 1 | 0.442 | 0.403 | 6 | 5 | 1 |
| A02 - A96 | 0.351 | 0.252 | 9 | 7 | 1 | 0.370 | 0.320 | 7 | 6 | 1 | 0.422 | 0.421 | 5 | 5 | 1 |
| Measures | Short axis length (pixels) | | | | | Orientation | | | | | Solidity | | | | |
| Comparison | $a$ | $c$ | $\hat{n}_\alpha$ | $n_\alpha$ | $\Theta_{\alpha,\gamma}$ | $a$ | $c$ | $\hat{n}_\alpha$ | $n_\alpha$ | $\Theta_{\alpha,\gamma}$ | $a$ | $c$ | $\hat{n}_\alpha$ | $n_\alpha$ | $\Theta_{\alpha,\gamma}$ |
| A02 - A03 | 0.267 | 0.005 | 284 | 330 | 1 | 0.265 | 0.000 | - | $2.38 \cdot 10^{18}$ | 0 | 0.273 | 0.013 | 124 | 126 | 1 |
| A02 - A09 | 0.263 | 0.000 | - | 3801 | 0 | 0.267 | 0.000 | - | $1.09 \cdot 10^{16}$ | 0 | 0.280 | 0.026 | 69 | 65 | 1 |
| A02 - A16 | 0.264 | 0.005 | 321 | 345 | 1 | 0.262 | 0.001 | - | $1.41 \cdot 10^{03}$ | 0 | 0.266 | 0.003 | 386 | 485 | 0 |
| A02 - A29 | 0.284 | 0.040 | 48 | 43 | 1 | 0.262 | 0.000 | - | $1.53 \cdot 10^{14}$ | 0 | 0.276 | 0.025 | 74 | 68 | 1 |
| A02 - A35 | 0.272 | 0.011 | 159 | 160 | 1 | 0.265 | 0.000 | - | $4.60 \cdot 10^{17}$ | 0 | 0.318 | 0.125 | 16 | 14 | 1 |
| A02 - A55 | 0.260 | 0.000 | - | 6616 | 0 | 0.265 | 0.000 | - | $1.65 \cdot 10^{20}$ | 0 | 0.267 | 0.005 | 267 | 319 | 1 |
| A02 - A65 | 0.291 | 0.057 | 34 | 30 | 1 | 0.262 | 0.001 | - | $3.20 \cdot 10^{03}$ | 0 | 0.275 | 0.015 | 117 | 112 | 1 |
| A02 - A85 | 0.311 | 0.103 | 20 | 17 | 1 | 0.265 | 0.000 | - | $8.27 \cdot 10^{16}$ | 0 | 0.321 | 0.177 | 11 | 10 | 1 |
| A02 - A96 | 0.295 | 0.069 | 27 | 25 | 1 | 0.261 | 0.000 | - | $1.13 \cdot 10^{16}$ | 0 | 0.298 | 0.077 | 26 | 23 | 1 |
| Measures | Equivalent diameter (pixels) | | | | | Aspect ratio | | | | | Circularity | | | | |
| Comparison | $a$ | $c$ | $\hat{n}_\alpha$ | $n_\alpha$ | $\Theta_{\alpha,\gamma}$ | $a$ | $c$ | $\hat{n}_\alpha$ | $n_\alpha$ | $\Theta_{\alpha,\gamma}$ | $a$ | $c$ | $\hat{n}_\alpha$ | $n_\alpha$ | $\Theta_{\alpha,\gamma}$ |
| A02 - A03 | 0.267 | 0.002 | 509 | 688 | 0 | 0.282 | 0.027 | 74 | 63 | 1 | 0.277 | 0.022 | 78 | 77 | 1 |
| A02 - A09 | 0.264 | 0.002 | - | 923 | 0 | 0.277 | 0.016 | 103 | 104 | 1 | 0.261 | 0.000 | - | $1.33 \cdot 10^{13}$ | 0 |
| A02 - A16 | 0.320 | 0.149 | 13 | 12 | 1 | 0.384 | 0.327 | 7 | 6 | 1 | 0.319 | 0.162 | 12 | 11 | 1 |
| A02 - A29 | 0.312 | 0.129 | 16 | 14 | 1 | 0.275 | 0.019 | 86 | 90 | 1 | 0.260 | 0.001 | - | 2313 | 0 |
| A02 - A35 | 0.311 | 0.106 | 19 | 17 | 1 | 0.281 | 0.042 | 44 | 40 | 1 | 0.259 | 0.000 | - | $1.22 \cdot 10^{15}$ | 0 |
| A02 - A55 | 0.313 | 0.112 | 18 | 16 | 1 | 0.318 | 0.154 | 13 | 12 | 1 | 0.289 | 0.056 | 36 | 31 | 1 |
| A02 - A65 | 0.326 | 0.137 | 16 | 13 | 1 | 0.306 | 0.094 | 20 | 19 | 1 | 0.313 | 0.113 | 18 | 16 | 1 |
| A02 - A85 | 0.330 | 0.167 | 13 | 11 | 1 | 0.290 | 0.068 | 27 | 25 | 1 | 0.338 | 0.238 | 9 | 8 | 1 |
| A02 - A96 | 0.351 | 0.252 | 9 | 7 | 1 | 0.326 | 0.149 | 13 | 12 | 1 | 0.337 | 0.187 | 11 | 10 | 1 |
| Measures | Roundness | | | | | | | | | | | | | | |
| Comparison | $a$ | $c$ | $\hat{n}_\alpha$ | $n_\alpha$ | $\Theta_{\alpha,\gamma}$ | | | | | | | | | | |
| A02 - A03 | 0.277 | 0.022 | 78 | 77 | 1 | | | | | | | | | | |
| A02 - A09 | 0.261 | 0.000 | - | $1.33 \cdot 10^{13}$ | 0 | | | | | | | | | | |
| A02 - A16 | 0.319 | 0.162 | 12 | 11 | 1 | | | | | | | | | | |
| A02 - A29 | 0.260 | 0.001 | - | 2313 | 0 | | | | | | | | | | |
| A02 - A35 | 0.259 | 0.000 | - | $1.2 \cdot 10^{15}$ | 0 | | | | | | | | | | |
| A02 - A55 | 0.289 | 0.056 | 36 | 31 | 1 | | | | | | | | | | |
| A02 - A65 | 0.313 | 0.113 | 18 | 16 | 1 | | | | | | | | | | |
| A02 - A85 | 0.338 | 0.238 | 9 | 8 | 1 | | | | | | | | | | |
| A02 - A96 | 0.337 | 0.187 | 11 | 10 | 1 | | | | | | | | | | |

**Table S6. Parameters of the exponential function $ae^{-cn}$ for cell nuclei morphology measures, theoretical minimum size ($n_\alpha$) and estimated one ($\hat{n}_\alpha$) for a $95\%$ ($\alpha = 0.05$) of statistical significance, and decision index $\Theta_{\alpha,\gamma}$ , for $\gamma = 5 \cdot 10^{-6}$.**

| $\gamma$ | $2.5 \cdot 10^{-6}$ | | $5 \cdot 10^{-6}$ | | $5 \cdot 10^{-5}$ | | $5 \cdot 10^{-4}$ | | |
|---|---|---|---|---|---|---|---|---|---|
| Comparison | $\Theta_{\alpha,\gamma}$ | $\delta_{\alpha,\gamma}$ | $\Theta_{\alpha,\gamma}$ | $\delta_{\alpha,\gamma}$ | $\Theta_{\alpha,\gamma}$ | $\delta_{\alpha,\gamma}$ | $\Theta_{\alpha,\gamma}$ | $\delta_{\alpha,\gamma}$ | $\hat{n}_\alpha$ |
| $\mathcal{N}(0,1)$ - $\mathcal{N}(0,1)$ | 0 | -490.25 | 0 | -490.25 | 0 | -490.25 | 0 | -490.25 | 0 |
| $\mathcal{N}(0,1)$ - $\mathcal{N}(0.01,1)$ | 0 | -3576 | 0 | -488.25 | 0 | -488.25 | 0 | -488.25 | 0 |
| $\mathcal{N}(0,1)$ - $\mathcal{N}(0.1,1)$ | 1 | 0.782 | 0 | -21.922 | 0 | -79.938 | 0 | -42.375 | 1237 |
| $\mathcal{N}(0,1)$ - $\mathcal{N}(0.25,1)$ | 1 | 9.523 | 1 | 5.848 | 0 | -5.840 | 0 | -12.945 | 186 |
| $\mathcal{N}(0,1)$ - $\mathcal{N}(0.5,1)$ | 1 | 0 | 1 | 0 | 1 | 0 | 1 | 0 | 45 |
| $\mathcal{N}(0,1)$ - $\mathcal{N}(0.75,1)$ | 1 | 0 | 1 | 0 | 1 | 0 | 1 | 0 | 22 |
| $\mathcal{N}(0,1)$ - $\mathcal{N}(1,1)$ | 1 | 0 | 1 | 0 | 1 | 0 | 1 | 0 | 0 |
| $\mathcal{N}(0,1)$ - $\mathcal{N}(1.5,1)$ | 1 | 0 | 1 | 0 | 1 | 0 | 1 | 0 | 0 |
| $\mathcal{N}(0,1)$ - $\mathcal{N}(2,1)$ | 1 | 0 | 1 | 0 | 1 | 0 | 1 | 0 | 0 |
| $\mathcal{N}(0,1)$ - $\mathcal{N}(2.5,1)$ | 1 | 0 | 1 | 0 | 1 | 0 | 1 | 0 | 0 |
| $\mathcal{N}(0,1)$ - $\mathcal{N}(3,1)$ | 1 | 0 | 1 | 0 | 1 | 0 | 1 | 0 | 0 |

**Table S7. Table of decision index $\Theta_{\alpha,\gamma}$, difference $\delta_{\alpha,\gamma} = A_{\alpha_\gamma} - A_{p(n_\gamma)}$ and estimated minimum size ($\hat{n}_\alpha$) for $\alpha = 0.05$ and $\gamma = 2.5 \cdot 10^{-6}, 5 \cdot 10^{-6}, 5 \cdot 10^{-5}, 5 \cdot 10^{-4}$.**

| | | $\mathcal{N}$ grid size | | | | | | | | $\mathcal{N}$ grid size | | | | | |
|---|---|---|---|---|---|---|---|---|---|---|---|---|---|---|---|
| Comparison | $\mathcal{F}$ reduction | $10$ | $20$ | $50$ | $100$ | $150$ | $200$ | Comparison | $\mathcal{F}$ reduction | $10$ | $20$ | $50$ | $100$ | $150$ | $200$ |
| $\mathcal{N}(0,1)$ - $\mathcal{N}(0,1)$ | 0.1 | 100 | 100 | 100 | 100 | 100 | 100 | $\mathcal{N}(0,1)$ - $\mathcal{N}(0.75,1)$ | 0.1 | 100 | 100 | 100 | 100 | 100 | 100 |
| | 0.2 | 100 | 100 | 100 | 100 | 100 | 100 | | 0.2 | 100 | 100 | 100 | 100 | 100 | 100 |
| | 1/3 | 100 | 100 | 100 | 100 | 100 | 100 | | 1/3 | 100 | 100 | 100 | 100 | 100 | 100 |
| | 0.5 | 100 | 100 | 100 | 100 | 100 | 100 | | 0.5 | 100 | 100 | 100 | 100 | 100 | 100 |
| | 1 | 100 | 100 | 100 | 100 | 100 | 100 | | 1 | 100 | 100 | 100 | 100 | 100 | 100 |
| $\mathcal{N}(0,1)$ - $\mathcal{N}(0.01,1)$ | 0.1 | 100 | 100 | 100 | 100 | 100 | 100 | $\mathcal{N}(0,1)$ - $\mathcal{N}(1,1)$ | 0.1 | 100 | 100 | 100 | 100 | 100 | 100 |
| | 0.2 | 100 | 100 | 100 | 100 | 100 | 100 | | 0.2 | 100 | 100 | 100 | 100 | 100 | 100 |
| | 1/3 | 100 | 100 | 100 | 100 | 100 | 100 | | 1/3 | 100 | 100 | 100 | 100 | 100 | 100 |
| | 0.5 | 100 | 100 | 100 | 100 | 100 | 100 | | 0.5 | 100 | 100 | 100 | 100 | 100 | 100 |
| | 1 | 100 | 100 | 100 | 100 | 100 | 100 | | 1 | 100 | 100 | 100 | 100 | 100 | 100 |
| $\mathcal{N}(0,1)$ - $\mathcal{N}(0.1,1)$ | 0.1 | 99 | 100 | 100 | 100 | 100 | 100 | $\mathcal{N}(0,1)$ - $\mathcal{N}(1.5,1)$ | 0.1 | 100 | 100 | 100 | 100 | 100 | 100 |
| | 0.2 | 100 | 100 | 100 | 100 | 100 | 100 | | 0.2 | 100 | 100 | 100 | 100 | 100 | 100 |
| | 1/3 | 100 | 100 | 100 | 100 | 100 | 100 | | 1/3 | 100 | 100 | 100 | 100 | 100 | 100 |
| | 0.5 | 100 | 100 | 100 | 100 | 100 | 100 | | 0.5 | 100 | 100 | 100 | 100 | 100 | 100 |
| | 1 | 100 | 100 | 100 | 100 | 100 | 100 | | 1 | 100 | 100 | 100 | 100 | 100 | 100 |
| $\mathcal{N}(0,1)$ - $\mathcal{N}(0.25,1)$ | 0.1 | 100 | 100 | 100 | 100 | 100 | 100 | $\mathcal{N}(0,1)$ - $\mathcal{N}(2,1)$ | 0.1 | 100 | 100 | 100 | 100 | 100 | 100 |
| | 0.2 | 100 | 100 | 100 | 100 | 100 | 100 | | 0.2 | 100 | 100 | 100 | 100 | 100 | 100 |
| | 1/3 | 100 | 100 | 100 | 100 | 100 | 100 | | 1/3 | 100 | 100 | 100 | 100 | 100 | 100 |
| | 0.5 | 100 | 100 | 100 | 100 | 100 | 100 | | 0.5 | 100 | 100 | 100 | 100 | 100 | 100 |
| | 1 | 100 | 100 | 100 | 100 | 100 | 100 | | 1 | 100 | 100 | 100 | 100 | 100 | 100 |
| $\mathcal{N}(0,1)$ - $\mathcal{N}(0.5,1)$ | 0.1 | 89 | 95 | 99 | 99 | 100 | 100 | $\mathcal{N}(0,1)$ - $\mathcal{N}(3,1)$ | 0.1 | 100 | 100 | 100 | 100 | 100 | 100 |
| | 0.2 | 96 | 99 | 100 | 100 | 100 | 100 | | 0.2 | 100 | 100 | 100 | 100 | 100 | 100 |
| | 1/3 | 100 | 99 | 100 | 100 | 100 | 100 | | 1/3 | 100 | 100 | 100 | 100 | 100 | 100 |
| | 0.5 | 99 | 100 | 100 | 100 | 100 | 100 | | 0.5 | 100 | 100 | 100 | 100 | 100 | 100 |
| | 1 | 100 | 100 | 100 | 100 | 100 | 100 | | 1 | 100 | 100 | 100 | 100 | 100 | 100 |

**Table S8. Table of results for different sizes of $\mathcal{N}$ and $\mathcal{F}$ grids. Each value represents the probability (%) of obtaining the same decision index $\Theta_{0.05,5}$ as the one shown in Table S9.**

| Variable | Cell body size | | | | | | | | |
|---|---|---|---|---|---|---|---|---|---|
| $\gamma$ | $2.5 \cdot 10^{-6}$ | | $5 \cdot 10^{-6}$ | | $5 \cdot 10^{-5}$ | | $5 \cdot 10^{-4}$ | | |
| Comparison | $\Theta_{\alpha,\gamma}$ | $\delta_{\alpha,\gamma}$ | $\Theta_{\alpha,\gamma}$ | $\delta_{\alpha,\gamma}$ | $\Theta_{\alpha,\gamma}$ | $\delta_{\alpha,\gamma}$ | $\Theta_{\alpha,\gamma}$ | $\delta_{\alpha,\gamma}$ | $\hat{n}_\alpha$ |
| Control - 1 nM Taxol | 1 | 8.67 | **0** | -4.12 | **0** | -41.26 | - | - | 670 |
| Control - 50 nM Taxol | 1 | 9.37 | 1 | 4.82 | **0** | -9.69 | - | - | 250 |
| 1 nM - 50 nM Taxol | 1 | 5.38 | 1 | 3.74 | **0** | -1.47 | **0** | 5.84 | 83 |
| Variable | Cell body perimeter | | | | | | | | |
| $\gamma$ | $2.5 \cdot 10^{-6}$ | | $5 \cdot 10^{-6}$ | | $5 \cdot 10^{-5}$ | | $5 \cdot 10^{-4}$ | | |
| Comparison | $\Theta_{\alpha,\gamma}$ | $\delta_{\alpha,\gamma}$ | $\Theta_{\alpha,\gamma}$ | $\delta_{\alpha,\gamma}$ | $\Theta_{\alpha,\gamma}$ | $\delta_{\alpha,\gamma}$ | $\Theta_{\alpha,\gamma}$ | $\delta_{\alpha,\gamma}$ | $\hat{n}_\alpha$ |
| Control - 1 nM Taxol | 1 | 1.37 | **0** | -17.84 | **0** | -69.41 | - | - | 1160 |
| Control - 50 nM Taxol | **0** | -5.75 | **0** | -29.94 | **0** | -90.28 | - | - | 1331 |
| 1 nM - 50 nM Taxol | 1 | 9.44 | 1 | 4.69 | **0** | -10.43 | - | - | 257 |
| Variable | Cell body roundness | | | | | | | | |
| $\gamma$ | $2.5 \cdot 10^{-6}$ | | $5 \cdot 10^{-6}$ | | $5 \cdot 10^{-5}$ | | $5 \cdot 10^{-4}$ | | |
| Comparison | $\Theta_{\alpha,\gamma}$ | $\delta_{\alpha,\gamma}$ | $\Theta_{\alpha,\gamma}$ | $\delta_{\alpha,\gamma}$ | $\Theta_{\alpha,\gamma}$ | $\delta_{\alpha,\gamma}$ | $\Theta_{\alpha,\gamma}$ | $\delta_{\alpha,\gamma}$ | $\hat{n}_\alpha$ |
| Control - 1 nM Taxol | 1 | 9.17 | **0** | -2.67 | **0** | -37.41 | - | - | 617 |
| Control - 50 nM Taxol | 1 | 3.42 | 1 | 2.57 | **0** | -0.25 | **0** | -2.87 | 47 |
| 1 nM - 50 nM Taxol | 1 | 2.39 | 1 | 1.84 | 1 | 0.05 | **0** | -1.59 | 29 |
| Variable | Protrusions binary | | | | | | | | |
| $\gamma$ | $2.5 \cdot 10^{-6}$ | | $5 \cdot 10^{-6}$ | | $5 \cdot 10^{-5}$ | | $5 \cdot 10^{-4}$ | | |
| Comparison | $\Theta_{\alpha,\gamma}$ | $\delta_{\alpha,\gamma}$ | $\Theta_{\alpha,\gamma}$ | $\delta_{\alpha,\gamma}$ | $\Theta_{\alpha,\gamma}$ | $\delta_{\alpha,\gamma}$ | $\Theta_{\alpha,\gamma}$ | $\delta_{\alpha,\gamma}$ | $\hat{n}_\alpha$ |
| Control - 1 nM Taxol | **0** | 0.00 | **0** | 0.00 | **0** | 0.00 | - | - | - |
| Control - 50 nM Taxol | 1 | 5.72 | 1 | 4.72 | 1 | 1.41 | **0** | -1.55 | 42 |
| 1 nM - 50 nM Taxol | 1 | 5.72 | 1 | 4.72 | 1 | 1.46 | **0** | -1.42 | 41 |
| Variable | Protrusions size | | | | | | | | |
| $\gamma$ | $2.5 \cdot 10^{-6}$ | | $5 \cdot 10^{-6}$ | | $5 \cdot 10^{-5}$ | | $5 \cdot 10^{-4}$ | | |
| Comparison | $\Theta_{\alpha,\gamma}$ | $\delta_{\alpha,\gamma}$ | $\Theta_{\alpha,\gamma}$ | $\delta_{\alpha,\gamma}$ | $\Theta_{\alpha,\gamma}$ | $\delta_{\alpha,\gamma}$ | $\Theta_{\alpha,\gamma}$ | $\delta_{\alpha,\gamma}$ | $\hat{n}_\alpha$ |
| Control - 1 nM Taxol | 1 | 11.99 | 1 | 1.15 | **0** | -31.16 | - | - | 563 |
| Control - 50 nM Taxol | 1 | 6.23 | 1 | 4.68 | **0** | -0.42 | **0** | -4.73 | 75 |
| 1 nM - 50 nM Taxol | 1 | 9.50 | 1 | 6.07 | **0** | -4.98 | **0** | -11.98 | 170 |
| Variable | Protrusions perimeter | | | | | | | | |
| $\gamma$ | $2.5 \cdot 10^{-6}$ | | $5 \cdot 10^{-6}$ | | $5 \cdot 10^{-5}$ | | $5 \cdot 10^{-4}$ | | |
| Comparison | $\Theta_{\alpha,\gamma}$ | $\delta_{\alpha,\gamma}$ | $\Theta_{\alpha,\gamma}$ | $\delta_{\alpha,\gamma}$ | $\Theta_{\alpha,\gamma}$ | $\delta_{\alpha,\gamma}$ | $\Theta_{\alpha,\gamma}$ | $\delta_{\alpha,\gamma}$ | $\hat{n}_\alpha$ |
| Control - 1 nM Taxol | 1 | 7.75 | **0** | -9.85 | **0** | -58.09 | - | - | 754 |
| Control - 50 nM Taxol | 1 | 5.54 | 1 | 4.29 | 1 | 0.15 | **0** | -3.40 | 58 |
| 1 nM - 50 nM Taxol | 1 | 6.77 | 1 | 4.77 | **0** | -1.63 | **0** | -6.71 | 98 |
| Variable | Protrusions length | | | | | | | | |
| $\gamma$ | $2.5 \cdot 10^{-6}$ | | $5 \cdot 10^{-6}$ | | $5 \cdot 10^{-5}$ | | $5 \cdot 10^{-4}$ | | |
| Comparison | $\Theta_{\alpha,\gamma}$ | $\delta_{\alpha,\gamma}$ | $\Theta_{\alpha,\gamma}$ | $\delta_{\alpha,\gamma}$ | $\Theta_{\alpha,\gamma}$ | $\delta_{\alpha,\gamma}$ | $\Theta_{\alpha,\gamma}$ | $\delta_{\alpha,\gamma}$ | $\hat{n}_\alpha$ |
| Control - 1 nM Taxol | **0** | -14.70 | **0** | -45.38 | **0** | -114.17 | - | - | 1695 |
| Control - 50 nM Taxol | 1 | 5.10 | 1 | 3.90 | **0** | -0.04 | **0** | -3.45 | 58 |
| 1 nM - 50 nM Taxol | 1 | 6.09 | 1 | 4.45 | **0** | -0.90 | **0** | -5.37 | 80 |
| Variable | Protrusions diameter | | | | | | | | |
| $\gamma$ | $2.5 \cdot 10^{-6}$ | | $5 \cdot 10^{-6}$ | | $5 \cdot 10^{-5}$ | | $5 \cdot 10^{-4}$ | | |
| Comparison | $\Theta_{\alpha,\gamma}$ | $\delta_{\alpha,\gamma}$ | $\Theta_{\alpha,\gamma}$ | $\delta_{\alpha,\gamma}$ | $\Theta_{\alpha,\gamma}$ | $\delta_{\alpha,\gamma}$ | $\Theta_{\alpha,\gamma}$ | $\delta_{\alpha,\gamma}$ | $\hat{n}_\alpha$ |
| Control - 1 nM Taxol | 1 | 9.64 | **0** | -4.80 | **0** | -45.90 | **0** | -12.24 | 707 |
| Control - 50 nM Taxol | 1 | 5.63 | 1 | 4.29 | **0** | -0.29 | **0** | -4.20 | 68 |
| 1 nM - 50 nM Taxol | 1 | 8.34 | 1 | 5.81 | **0** | -2.49 | **0** | -8.57 | 127 |

**Table S9. Table of decision index $\Theta_{\alpha,\gamma}$, difference $\delta_{\alpha,\gamma} = A_{\alpha_\gamma} - A_{p(n_\gamma)}$ and estimated minimum size $\hat{n}_\alpha$ for $\alpha = 0.05$ and $\gamma = 2.5 \cdot 10^{-6}, 5 \cdot 10^{-6}, 5 \cdot 10^{-5}, 5 \cdot 10^{-4}$.**

| Variables | | Cell body size | | | | Cell body perimeter | | | | Cell body roundness | | | | Protrusions binary | | | |
|---|---|---|---|---|---|---|---|---|---|---|---|---|---|---|---|---|---|
| | | $\mathcal{N}$ grid size | | | | $\mathcal{N}$ grid size | | | | $\mathcal{N}$ grid size | | | | $\mathcal{N}$ grid size | | | |
| Comparison | reduction | 10 | 20 | 50 | 100 | 10 | 20 | 50 | 100 | 10 | 20 | 50 | 100 | 10 | 20 | 50 | 100 |
| Control - $1$ nM Taxol | 0.01 | 80 | 98 | 82 | 94 | 95 | 100 | 99 | 100 | 71 | 92 | 71 | 79 | 100 | 100 | 100 | 100 |
| | 0.02 | 86 | 99 | 79 | 99 | 99 | 100 | 100 | 100 | 71 | 98 | 84 | 95 | 100 | 100 | 100 | 100 |
| | 0.1 | 100 | 100 | 99 | 100 | 100 | 100 | 100 | 100 | 93 | 100 | 98 | 100 | 100 | 100 | 100 | 100 |
| | 0.2 | 100 | 100 | 100 | 100 | 100 | 100 | 100 | 100 | 96 | 100 | 100 | 100 | 100 | 100 | 100 | 100 |
| | 1/3 | 100 | 100 | 100 | 100 | 100 | 100 | 100 | 100 | 100 | 100 | 100 | 100 | 100 | 100 | 100 | 100 |
| | 0.5 | 100 | 100 | 100 | 100 | 100 | 100 | 100 | 100 | 100 | 100 | 100 | 100 | 100 | 100 | 100 | 100 |
| Control - $50$ nM Taxol | 0.01 | 93 | 100 | 99 | 100 | 100 | 100 | 100 | 100 | 89 | 100 | 93 | 94 | 92 | 99 | 88 | 87 |
| | 0.02 | 100 | 100 | 100 | 100 | 100 | 100 | 100 | 100 | 100 | 100 | 96 | 100 | 94 | 99 | 87 | 98 |
| | 0.1 | 100 | 100 | 100 | 100 | 100 | 100 | 100 | 100 | 100 | 100 | 100 | 100 | 100 | 100 | 100 | 100 |
| | 0.2 | 100 | 100 | 100 | 100 | 100 | 100 | 100 | 100 | 100 | 100 | 100 | 100 | 100 | 100 | 100 | 100 |
| | 1/3 | 100 | 100 | 100 | 100 | 100 | 100 | 100 | 100 | 100 | 100 | 100 | 100 | 100 | 100 | 100 | 100 |
| | 0.5 | 100 | 100 | 100 | 100 | 100 | 100 | 100 | 100 | 100 | 100 | 100 | 100 | 100 | 100 | 100 | 100 |
| $1$ nM - $50$ nM Taxol | 0.01 | 95 | 99 | 98 | 98 | 98 | 100 | 100 | 100 | 80 | 94 | 81 | 89 | 79 | 98 | 78 | 91 |
| | 0.02 | 98 | 100 | 100 | 100 | 99 | 100 | 100 | 100 | 89 | 99 | 93 | 96 | 90 | 99 | 90 | 99 |
| | 0.1 | 100 | 100 | 100 | 100 | 100 | 100 | 100 | 100 | 100 | 100 | 100 | 100 | 100 | 100 | 98 | 100 |
| | 0.2 | 100 | 100 | 100 | 100 | 100 | 100 | 100 | 100 | 100 | 100 | 100 | 100 | 100 | 100 | 100 | 100 |
| | 1/3 | 100 | 100 | 100 | 100 | 100 | 100 | 100 | 100 | 100 | 100 | 100 | 100 | 100 | 100 | 100 | 100 |
| | 0.5 | 100 | 100 | 100 | 100 | 100 | 100 | 100 | 100 | 100 | 100 | 100 | 100 | 100 | 100 | 100 | 100 |
| Variables | | Protrusions size | | | | Protrusions perimeter | | | | Protrusions length | | | | Protrusions diameter | | | |
| | | $n$ grid size | | | | $n$ grid size | | | | $n$ grid size | | | | $n$ grid size | | | |
| Comparison | reduction | 10 | 20 | 50 | 100 | 10 | 20 | 50 | 100 | 10 | 20 | 50 | | 10 | 20 | 50 | 100 |
| Control - $1$ nM Taxol | 0.01 | 45 | 28 | 46 | 51 | 76 | 79 | 80 | 82 | 88 | 84 | 97 | 98 | 76 | 84 | 76 | 77 |
| | 0.02 | 36 | 35 | 50 | 60 | 74 | 65 | 79 | 90 | 86 | 89 | 99 | 100 | 78 | 79 | 85 | 90 |
| | 0.1 | 51 | 36 | 61 | 71 | 78 | 58 | 98 | 100 | 98 | 98 | 100 | 100 | 86 | 82 | 97 | 99 |
| | 0.2 | 64 | 24 | 59 | 83 | 89 | 57 | 100 | 100 | 99 | 100 | 100 | 100 | 91 | 95 | 100 | 100 |
| | 1/3 | 68 | 22 | 75 | 97 | 97 | 68 | 100 | 100 | 100 | 100 | 100 | 100 | 98 | 98 | 100 | 100 |
| | 0.5 | 66 | 17 | 82 | 99 | 98 | 75 | 100 | 100 | 100 | 100 | 100 | 100 | 100 | 100 | 100 | 100 |
| Control - $50$ nM Taxol | 0.01 | 76 | 79 | 91 | 95 | 82 | 73 | 84 | 99 | 74 | 69 | 88 | 93 | 74 | 67 | 90 | 97 |
| | 0.02 | 91 | 98 | 99 | 99 | 85 | 85 | 95 | 98 | 75 | 89 | 93 | 98 | 83 | 86 | 94 | 98 |
| | 0.1 | 100 | 100 | 100 | 100 | 100 | 100 | 100 | 100 | 100 | 100 | 100 | 100 | 100 | 100 | 100 | 100 |
| | 0.2 | 100 | 100 | 100 | 100 | 100 | 100 | 100 | 100 | 100 | 100 | 100 | 100 | 100 | 100 | 100 | 100 |
| | 1/3 | 100 | 100 | 100 | 100 | 100 | 100 | 100 | 100 | 100 | 100 | 100 | 100 | 100 | 100 | 100 | 100 |
| | 0.5 | 100 | 100 | 100 | 100 | 100 | 100 | 100 | 100 | 100 | 100 | 100 | 100 | 100 | 100 | 100 | 100 |
| $1$ nM - $50$ nM Taxol | 0.01 | 71 | 67 | 96 | 99 | 73 | 67 | 92 | 100 | 72 | 75 | 90 | 95 | 75 | 72 | 94 | 100 |
| | 0.02 | 92 | 80 | 100 | 100 | 87 | 84 | 99 | 100 | 94 | 90 | 98 | 100 | 86 | 87 | 99 | 100 |
| | 0.1 | 100 | 100 | 100 | 100 | 98 | 100 | 100 | 100 | 99 | 100 | 100 | 100 | 100 | 100 | 100 | 100 |
| | 0.2 | 100 | 100 | 100 | 100 | 100 | 100 | 100 | 100 | 100 | 100 | 100 | 100 | 100 | 100 | 100 | 100 |
| | 1/3 | 100 | 100 | 100 | 100 | 100 | 100 | 100 | 100 | 100 | 100 | 100 | 100 | 100 | 100 | 100 | 100 |
| | 0.5 | 100 | 100 | 100 | 100 | 100 | 100 | 100 | 100 | 100 | 100 | 100 | 100 | 100 | 100 | 100 | 100 |

**Table S10. Table of results for different sizes of $\mathcal{N}$ and $\mathcal{F}$ grids. Each value represents the probability (%) of obtaining the same decision index $\Theta_{0.05,5e-06}$ as the one shown in Table S9.**
